# Identifying Lineage-specific Targets of Darwinian Selection by a Bayesian Analysis of Genomic Polymorphisms and Divergence from Multiple Species

**DOI:** 10.1101/367482

**Authors:** Shilei Zhao, Tao Zhang, Qi Liu, Yongming Liu, Hao Wu, Bing Su, Peng Shi, Hua Chen

## Abstract

We present a method that jointly analyzes the polymorphism and divergence sites in genomic sequences of multiple species to identify the genes under positive or negative selection and pinpoints the occurrence time of selection to a specific lineage of the species phylogeny. This method integrates population genetics models using the Bayesian Poisson random field framework and combines information over all gene loci to boost the power to detect selection. The method provides posterior distributions of the fitness effects of each gene along with parameters associated with the evolutionary history, including the species divergence times and effective population sizes of external species. A simulation is performed, and the results demonstrate that our method provides accurate estimates of these population genetic parameters.

The proposed method is applied to genomic sequences of humans, chimpanzees, gorillas and orangutans, and a spatial and temporal map is constructed of the natural selection that occurred during the evolutionary history of the four Hominidae species. In addition to FOXP2 and other known genes, we identify a new list of lineage-specific targets of Darwinian selection. The positively selected genes in the human lineage are enriched in pathways of gene expression regulation, immune system, metabolism etc. Interestingly, some pathways, such as gene expression, are significantly enriched with positively selected genes, whereas other pathways, such as metabolism, are enriched with both positively and negatively selected genes. Our analysis provides insights into Darwinian evolution in the coding regions of humans and great apes and thus serves as a basis for further molecular and functional studies.

## 1. Introduction

Comparing the genomic sequences of multiple species is useful for studying evolutionary mechanism and identifying functionally important genes (Fay et al., 2002; Smith and Eyre-Walker, 2002; Clark et al., 2003; Rogers and Gibbs, 2014). Codon comparison across different species is perhaps the most widely used approach in comparative genomics (Li et al., 1985; Hughes and Nei, 1988; Yang, 1998). This method estimates the ratio of the number of replacement sites to the number of synonymous sites for different species (*dN/dS* ratio) to detect the genes under recurrent directional selection. The method was further developed to identify ratio changes on a specific branch along the phylogenetic tree (Yang, 1998; Yang and Nielsen, 2002; Yang, 2007; Zhang et al., 2005). These methods were designed to analyze single gene locus with one sequence per species without making use of population genetic samples, and they have been extensively used to identify targets of natural selection in various species (e.g., Clark et al. (2003)).

The McDonald-Kreitman test (MK test) is another method that uses codon replacement to detect Darwinian selection. The MK test compares coding sequences from two species and requires that at least one of the two species has multiple sequences. Polymorphism sites (within each species) and divergence sites (between species) of coding sequences are identified in the sample and further classified as synonymous or replacement (non-synonymous) sites. A contingency table is constructed based on the above four site types, and the chi-square test is used to examine the equality of the within-species ratios and the between-species ratios of replacements over synonymous sites. The underlying assumptions of the MK test are that (1) synonymous sites are under neutrality while non-synonymous sites are potentially under positive or negative selection; and (2) if one of the two species is under long-term recurrent selection since their divergence, then the ratio of the non-synonymous site number over the synonymous site number for the between-species divergence sites will be significantly greater than that of the within-species sites, and this situation is reversed for negative selection (McDonald and Kreitman, 1991).

Comparing within-species polymorphism to between-species divergence improves the power of detecting selection and enables the ability to control for the inter-locus heterogeneity of mutation rates, and it can also distinguish between positive selection and the relaxation of negative selection (Wyckoff et al., 2000). However, the MK test analyzes each gene individually and has limited power to detect loci under weak or moderate selection, especially when the numbers of these sites are small. Moreover, the chi-square test is not model based and cannot provide the strength and direction of selection acting on mutations in the genome. To tackle these problems, Bustamante et al. (2001) extended the MK test to embrace the population genetic model using the Poisson random field theory developed by Sawyer and Hartl (1992). The McDonald-Kreitman Poisson random field method (MKPRF) created by Bustamante et al. (2001) assumes that the number of sites in the MK table follows Poisson distributions, and their means are parameterized as functions of the population history, mutation rates and selection intensity. The method then uses a Bayesian approach to combine information across different gene loci and obtain the posterior distributions of selection intensity for non-synonymous sites of each gene.

Bustamante et al. (2005) applied the MKPRF Bayesian approach to analyze 11, 624 coding regions of 39 human and 1 chimpanzee sequences, and they identified 304 genes under rapid evolution since the divergence of the two species. However, the MKPRF approach applies only to directionless comparisons of two species; therefore, the method cannot determine whether selection occurred in the human or the chimpanzee lineage.

With the development of sequencing technology, abundant population genomic data are now available for multiple species. However, few efficient methods are available that can simultaneously analyze genomic polymorphisms and divergence from multiple species. The method developed in this paper is designed to satisfy a growing request for such methods, and it is an extension of the MKPRF method that simultaneously analyzes polymorphism sites and divergence sites from multiple species (aka, high-dimensional MKPRF, HDMKPRF).

Based on the joint pattern of divergence and polymorphism sites from multiple species, the proposed method not only identifies genes under selection in any of the analyzed species but also pinpoints the occurrence time of selection to a specific lineage of the species phylogeny. Compared with codon-based phylogenetic methods, such as PAML (phylogenetic analysis by maximum likelihood), the new method gains power by jointly analyzing both between-species divergence and within-species polymorphism sites and combining the information from all gene loci with a Bayesian approach. Furthermore, the method provides estimates of phylogenetic and population genetic parameters, such as the divergence times of species, mutation rates of each gene loci and effective population sizes of different species. The method constructs the spatial and temporal landscape of natural selection during the evolutionary history of the species. We applied the method to the genomic sequences of four species of Hominidae: humans, chimpanzees, gorillas and orangutans. We identified a set of genes under rapid adaptation in the human and chimpanzee lineages as well as genes under weak purifying selection. The positively selected genes are enriched in pathways related to gene expression regulation, immunity, metabolism, neurological diseases, etc, thereby providing a natural selection map for further investigation of the molecular mechanism in Hominidae evolution.

## Methods

### Poisson random field model (PRF) for three species

In the scenario involving three species (Figure 1), we analyze *n*_1_, *n*_2_ and *n*_3_ aligned sequences for a coding region from the three species. If we assume an infinite-sites mutation model with no introgression among species, the polymorphism and divergence sites can be classified into 14 types according to their joint patterns across the three species (see Table 1). For example, polymorphism sites that are segregating within species 1 and fixed in species 2 and 3 are denoted as *P*_1∼(2,3)_; divergence sites with one allele that is fixed in species 1 and another that is fixed in species 2 and 3 are denoted as *D*_1∼(2,3)_ etc. Following the MK test, a chi-square test with six degrees of freedom can be immediately constructed based on Table 1 to test the deviation of entries of the contingency table from randomness. In the following sections, we set up the model using the more efficient Bayesian Poisson random field framework.

**Figure 1:**
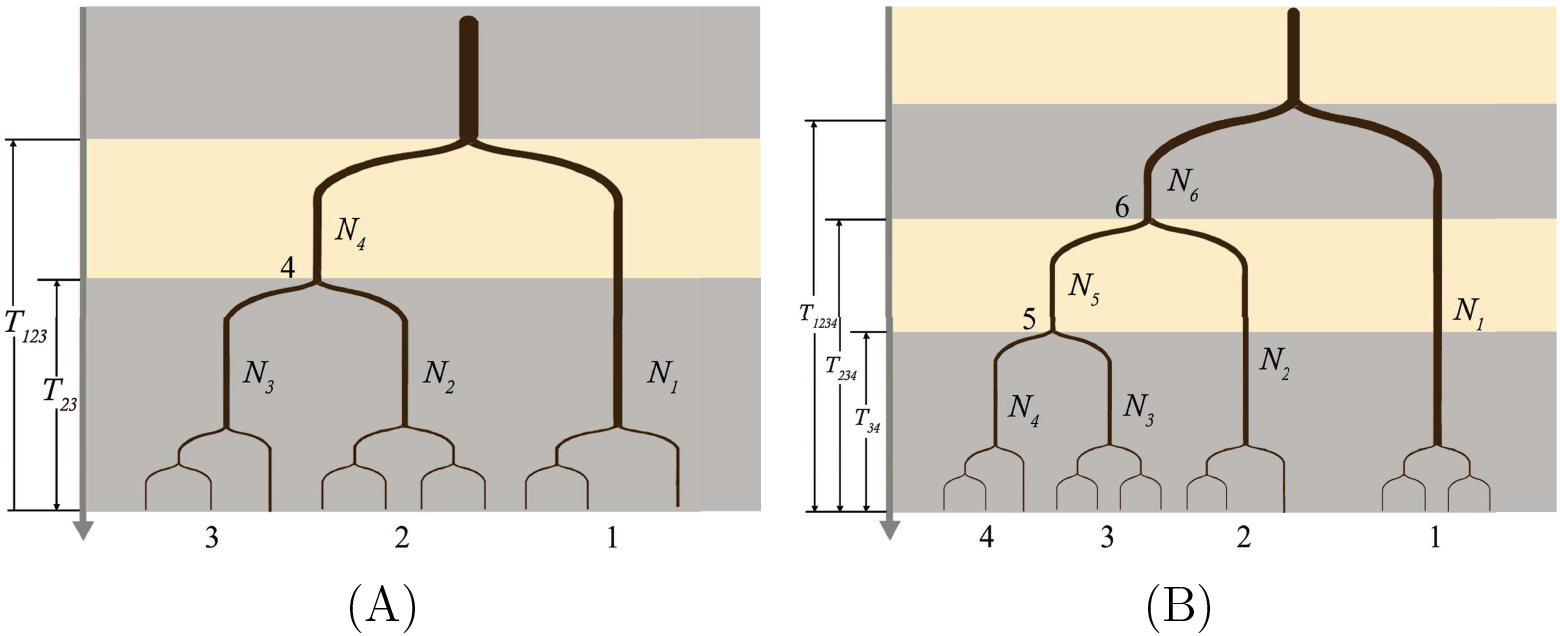
An illustration of the genealogies for three species and four species and the parameters.

**Table 1:** Three-species McDonald-Kreitman table.

|  |  | Synonymous | Replacement |
| --- | --- | --- | --- |
| $1 \sim (2, 3)$ | Divergence $D_{1 \sim (2,3)}$<br>Polymorphism $P_{1 \sim (2,3)}$ | $\theta_{s,1}(T_{12} - \frac{1}{2}T_{23} + M(n_1))$<br>$\theta_{s,1}L(n_1)$ | $\theta_{r,1}\frac{2\gamma_1}{1-e^{(-2\gamma_1)}}(T_{12} - \frac{1}{2}T_{23} + G(n_1, \gamma_1))$<br>$\theta_{r,1}\frac{2\gamma_1}{1-e^{(-2\gamma_1)}}F(n_1, \gamma_1)$ |
| $2 \sim (1, 3)$ | Divergence $D_{2 \sim (1,3)}$<br>Polymorphism $P_{2 \sim (1,3)}$ | $\theta_{s,2}(\frac{1}{2}v_3T_{23} + M(n_2))$<br>$\theta_{s,2}L(n_2)$ | $\theta_{r,2}\frac{2\gamma_2}{1-e^{(-2\gamma_2)}}(\frac{1}{2}v_2T_{23} + G(n_2, \gamma_2))$<br>$\theta_{r,2}\frac{2\gamma_2}{1-e^{(-2\gamma_2)}}F(n_2, \gamma_2)$ |
| $3 \sim (1, 2)$ | Divergence $D_{3 \sim (1,2)}$<br>Polymorphism $P_{3 \sim (1,2)}$ | $\theta_{s,3}(\frac{1}{2}v_3T_{23} + M(n_3))$<br>$\theta_{s,3}L(n_3)$ | $\theta_{r,3}\frac{2\gamma_3}{1-e^{(-2\gamma_3)}}(\frac{1}{2}v_3T_{23} + G(n_3, \gamma_3))$<br>$\theta_{r,3}\frac{2\gamma_3}{1-e^{(-2\gamma_3)}}F(n_3, \gamma_3)$ |
| $(2, 3) \sim 1$ | Polymorphism $P_{(2,3) \sim 1}$ | $\theta_{s,4}H(n_2, n_3, N_2, N_3, T_{23})$ | $\theta_{r,4}\frac{2\gamma_4}{1-e^{(-2\gamma_4)}}I(n_2, n_3, \gamma_2, \gamma_3, \gamma_4)$ |

In the Poisson random field modeling assumption, each of the 14 entries in Table 1 follows a Poisson distribution, and the means are parameterized with population genetic models (Sawyer and Hartl, 1992). The population genetic parameters **Γ** include: the divergence time between species 1 and the common ancestor of species 2 and 3 (*T*_123_), the divergence time between species 2 and 3 (*T*_23_), and the effective haploid population sizes of the three species (*N*_1_, *N*_2_, and *N*_3_). For each gene locus *i*, mutation rate 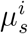 is observed for synonymous sites, mutation rate 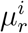 is observed for replacement sites, and selection intensities 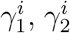 and 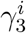 are observed in the three species.

Synonymous polymorphism sites in species 3, which are denoted by 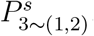, are neutral mutations that have occurred in species 3 since the divergence of species 2 and 3. If the divergence time is sufficiently large, then the population allele frequency *x* of 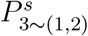 follows a stationary distribution (Sawyer and Hartl, 1992) 
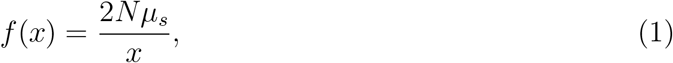
where *N* = *N*_3_. Thus, the expected number in a sample with *n*_3_ sequences is: 
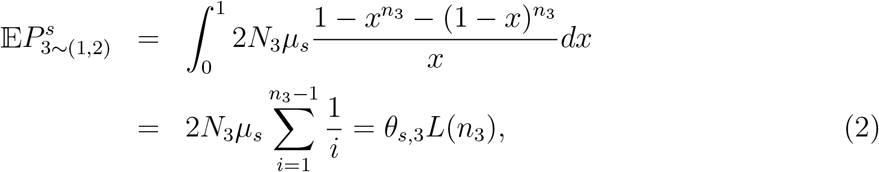
with 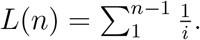.

The synonymous divergence sites, 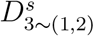, include those sites segregating in the population but fixed in the sample: 
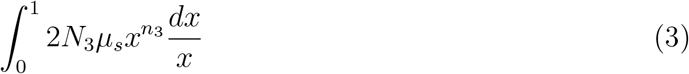

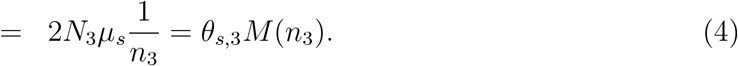

The synonymous divergence sites also include those sites fixed in the population of species 3: *μ_s_T_23_*. If we scale time *T*_23_ in units of *N*_1_, then 
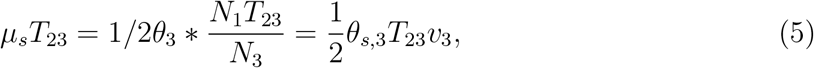
and 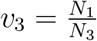.

For non-synonymous (replacement) sites, we know that the stationary population allele frequency under selection is as follows: 
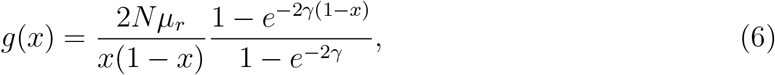
where *x* is the population allele frequency and γ = *Ns*, with *s* representing the selection intensity (Sawyer and Hartl, 1992). 1 The fixation rate is: 
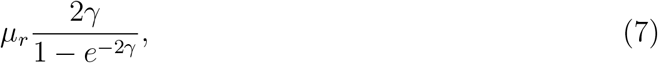

Thus, the number of replacement sites along branch leading to species 3 is: 
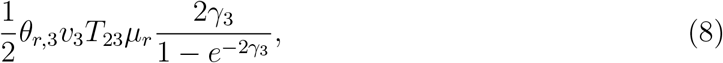
where *T*_23_ is in units of *N*_1_ generations.

The number of replacement sites fixed in the sample *n*_3_ is as follows: 
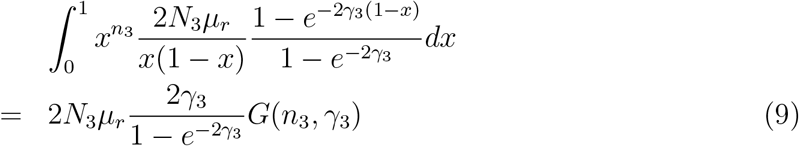
with 
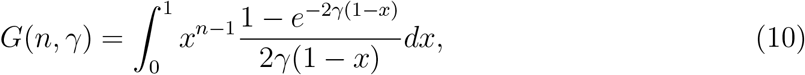

Similarly, the expected number of segregating replacement sites, 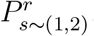, is: 
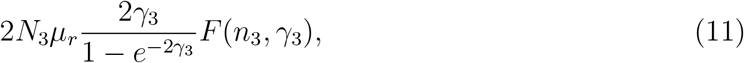
with 
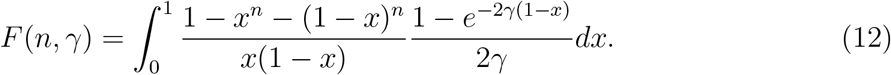

Through similar logic, we can obtain the expected number of divergence and polymorphism sites for *P*_2∼(1,3)_, *D*_2∼(1,3)_, *P*_1∼(2,3)_ and *D*_1∼(2,3)_ (see Table 1 for details).

In the aforementioned paragraphs, we assume that the divergence time *T*_23_ is sufficiently large; thus, the chance of observing polymorphism sites shared by species 2 and 3, *P*_(2,3)∼1_, has a very low probability. However, for closely-related species with *T*_23_ ≤ TMRCA of *n*_2_, the expected number of neutral polymorphic sites segregating in species 2 and 3, *P*_(2,3)∼1_, cannot be ignored and should be calculated from the joint allele frequency of species 2 and 3 *f* (*y*, *z*|*T*_23_, *N*_2_, *N*_3_) (see AppendixA for details): 
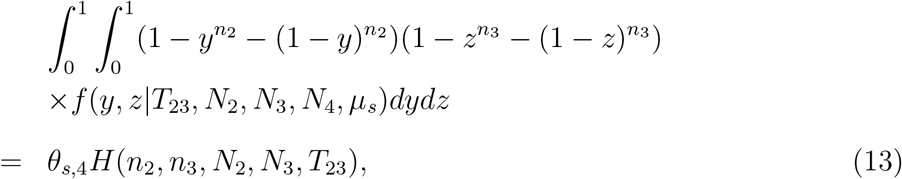
where *θ_s,4_* = 2*N*_4_*μ_s_* stands for scaled mutation rate in species 4 and *y* and *z* are the allele frequencies in species 2 and 3, respectively. Additionally, 
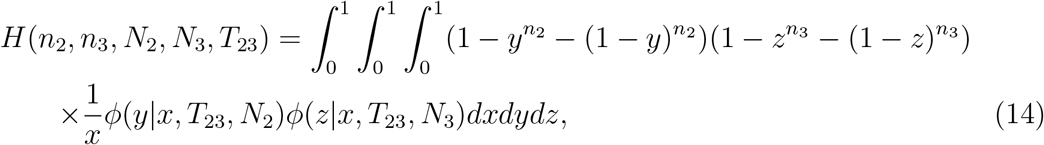
with *ϕ*(*y*|*x*, *T*, *N*) representing the transient allele frequency distribution conditional on an initial frequency *x*, population size *N* and time *T* (see Equation 25 for the detailed form). The expected number of selected polymorphic sites segregating in species 2 and 3 is 
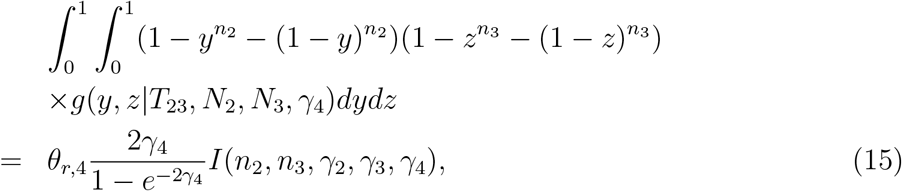
where *γ*_2_, *γ*_3_, and *γ*_4_ are the selection coefficients in species 2, 3 and, 4 respectively. And, 
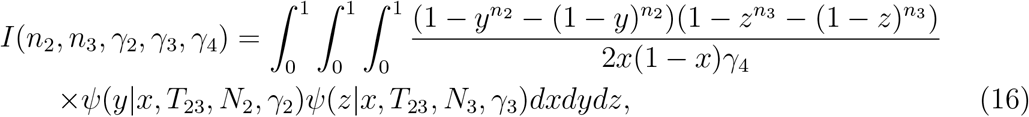
with *ψ* (*y*|*x*, *T*, *N*, *γ*) representing the transient allele frequency distribution conditional on an initial frequency *x*, population size *N*, time *T* and selection intensity *γ* (see Appendix A for details).

Accordingly, when under the assumption *T*_23_ ≤ TMRCA of *n*_2_, 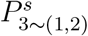 and other entries in Table 1 should be derived in a new form, which can be found in Appendix A.

Assuming a Poisson random field model, the joint probability of the data given parameter 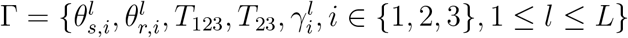 is the product of the individual entries of Table 1 for all the *L* gene loci: 
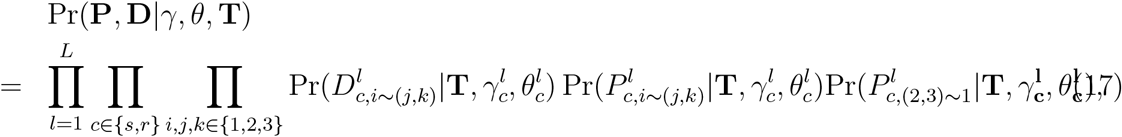
where Pr(·) denotes the Poisson distributions.

### Poisson random field model (PRF) for four species

The phylogeny of four species is shown in Figure 1 (B). Similar to the three-species case, the four-species data contains branch-specific divergence and polymorphism sites (e.g., *D*_1∼(2,3,4)_, *P*_1∼(2,3,4)_, etc.). In addition, there are some unique site patterns corresponding to internal branches connecting species 5 and 6. Mutations that occur on this branch could have generated multiple site patterns in the modern four-species samples. These patterns could be created by polymorphism sites shared by species 3 and 4, which is denoted by *P*(_3,4_)∼(_1,2_); by sites with one allele type fixed in species 3 and 4, and the other allele type fixed in species 1 and 2, which is denoted by *D*(_3,4_)∼(_1,2_); or by sites with one allele type that is fixed in species 3 and another fixed in species 1, 2 and 4 (or perhaps the reverse). Since we assume no migration or introgression between species and an infinite-sites mutation model, common mutations shared between species 3 and 4 can only be descended from ancestral mutations existing in *N*_5_. Similar to *P*_s_,_(2,3)∼1_ in the three-species scenario, the expected number of neutral polymorphic sites segregating in species 3 and 4 is as follows: 
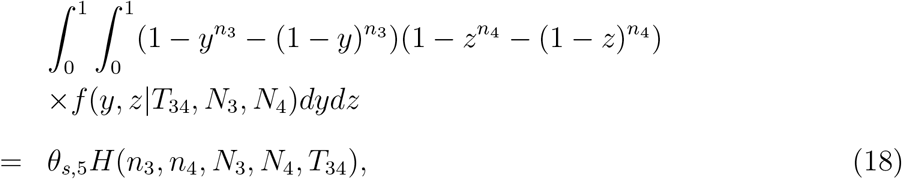
where *y* and *z* now represent the allele frequencies in species 3 and 4, respectively (see Appendix A for details). Similarly, the expected number of polymorphic replacement sites segregating in species 3 and 4 is 
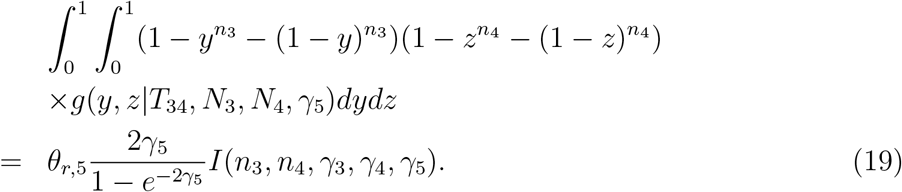

If the divergence time between species 3 and 4 is sufficiently large, then these common ancestral polymorphic sites are mostly lost or fixed, the values of *H*(*n*_3_, *n*_4_) and *I*(*n*_3_, *n*_4_) become negligible, and *P*_(3,4)∼(1,2)_ collapses into *P*_3∼(1,2,4)_, *P*_4∼(1,2,3)_ and *D*_(3,4)∼(1,2)_. Thus, although *P*_(3,4)∼(1,2)_ exists, the number could be only represent a small proportion and provides limited information for inference. This is consistent with the genomic data we observed in the four Hominidae species (see the result section), with *P*_(n_3_, n_4_)∼(1,2)_ only representing 0.2896% of the total number of segregating sites (197, 878).

The expected number of fixed synonymous sites 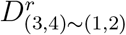 includes two components: 
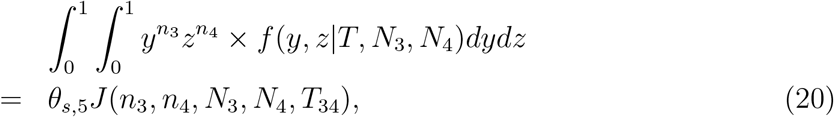
and 
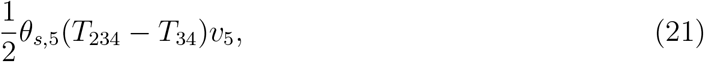
with υ_5_ = *N*_1_/*N*_5_.

Similarly, the expected number of fixed non-synonymous sites 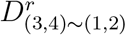 includes two parts: 
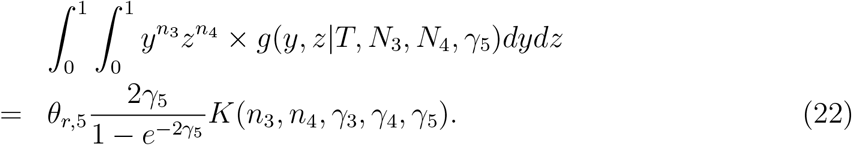
and 
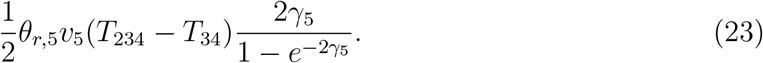

The first part contains a small proportion when the divergence time *T*_34_ is large.

## Results

### Simulation

We tested the performance of the method using simulation data. We simulated 20, 000 genes for the four-species scenario in Figure 1, and the population history parameters were set to approximate the evolutionary history of humans, chimpanzees, gorillas and orangutans inferred in previous studies (Prado-Martinez et al., 2013), and the sample includes *n*_4_ = 10, *n*_3_ = 24, *n*_2_ = 20, and *n*_1_ = 20 haplotypes from the four species, respectively. We set the effective haploid population sizes at *N*_2_ = *N*_1_, *N*_3_ = 1.2*N*_1_, and *N*_4_ = 0.8*N*_1_. The divergence times were *T*_34_ = 4, *T*_234_ = 6 and *T*_1234_ = 12 in units of 2*N*_1_. The scaled mutation rate for synonymous sites for each gene locus, *θ_s_* = 2*N*_1*μ_s_*_, and the scaled mutation rate for replacement sites, *θ_r_* = 2*N*_1*μ_r_*_, were chosen from several fixed values 1,2, …, 5. Among the 20, 000 genes, 1400 were under selection in species 4 or 5 (the common ancestor of humans and chimpanzees), and the other 18, 600 genes were neutral. The selection intensities *γ*_*i*_, *i* = 4, 5 of every 100 genes were chosen from fixed values in the range of (−6, −4, −2, 0, 2, 4, 6). Given the values of these parameters, simulated data were generated from Poisson distributions, and the means were calculated based on the formulae in Table 2. We then applied the method to the simulated data, and the maximum a posteriori (MAP) estimates of the parameters were recorded. The above simulations were repeated for 100 times. The boxplots of the inferred selection intensity, scaled mutation rates and divergence times are shown in Figure 2. For the global parameters, such as the divergence times *T*_1234_, *T*_234_ and *T*_34_, the inferred values are accurate and unbiased since sufficient information for these parameters is derived from the Bayesian joint analysis of all 20, 000 gene loci. The other locus-specific parameters, including the selection intensity and mutation rates of different branches, are generally also unbiased. However, we note that for selection intensity, when under strong negative selection, the inferred values become biased toward smaller values, because few or zero divergence and polymorphism sites are observed when the gene is under strong negative selection, which provides limited information for parameter inference (Bustamante et al., 2005). Overall, we evaluated the performance of HDMKPRF with simulated data, and the boxplots shown in Figure 2 demonstrate that the inferred parameter values well match the true values.

**Figure 2:**
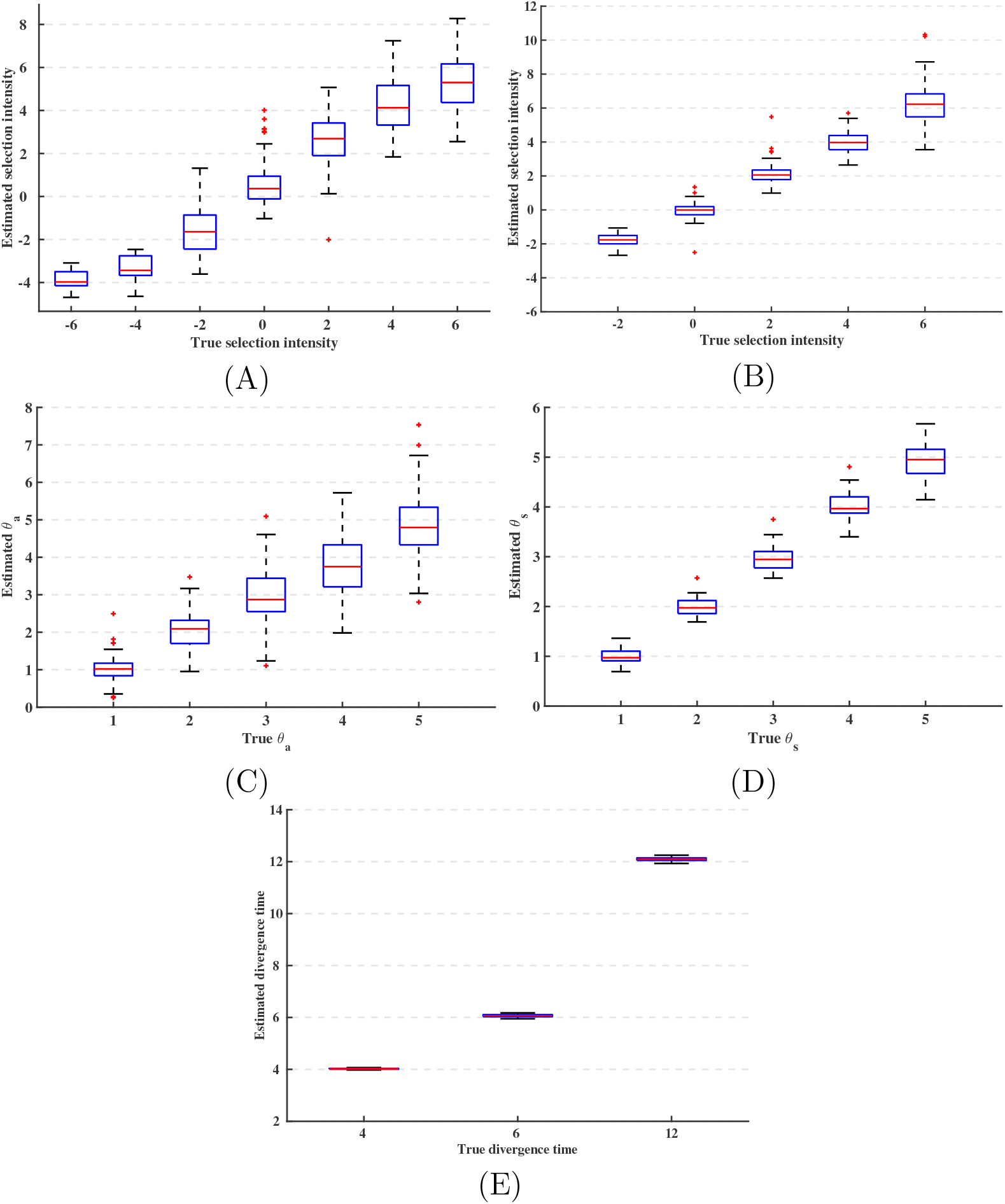
Comparison of the true and inferred selection coefficients for the four species in the simulation of 20,000 gene loci: (A) selection coefficient *γ*_4_; (B)selection coefficient *γ*_5_; (C) scaled mutation rate; (D) scaled synonymous mutation rate; and (E) divergence times *T*_34_, *T*_234_ and *T*_1234_.

### Selection in Hominidae

We applied the HDMKPRF method to multiple genomic sequences of humans, chimpanzees, gorillas and orangutans from Prado-Martinez et al. (2013). The data were generated via NGS technology with an average sequencing depth of 25, and details of the SNP calling pipeline and filtering criteria can be found in the original paper (Prado-Martinez et al., 2013). After excluding several individuals based on further criteria described in Cagan et al. (2016), the final dataset in our analysis includes *Pongo pygmaeus* (5), *Gorrilla gorilla* (12), *Pan troglodytes ellioti* (9) and *Homo sapiens* (9). We aligned the sequences of 23, 362 genes for these samples, from which 5, 429 genes with no protein coding information were excluded from the analysis. For the remaining 17, 933 genes (17, 234 from autosomes and 699 from X chromosome), we used ANNOVAR (Wang et al., 2010) to annotate the SNPs in the coding regions. Synonymous and replacement polymorphism and divergence sites were identified to construct the four-species MK tables for each gene (see entries of Table 2). In total, 133 genes with no divergence and segregating sites in all lineages were excluded from the four-species MK tables.

**Table 2:** Four-species McDonald-Kreitman table.

|  |  | Synonymous | Replacement |
| --- | --- | --- | --- |
| $1 \sim (2, 3, 4)$ | Divergence $D_{1 \sim (2,3,4)}$ | $\theta_{s,1}(T_{1234} - \frac{1}{2}T_{234} + M(n_1))$ | $\theta_{r,1} \frac{2\gamma_1}{1-e^{-2\gamma_1}}(T_{1234} - \frac{1}{2}T_{234} + G(n_1, \gamma_1))$ |
| | Polymorphism $P_{1 \sim (2,3,4)}$ | $\theta_{s,1}L(n_1)$ | $\theta_{r,1} \frac{2\gamma_1}{1-e^{-2\gamma_1}}F(n_1, \gamma_1)$ |
| $2 \sim (1, 3, 4)$ | Divergence $D_{2 \sim (1,3,4)}$ | $\theta_{s,2}(\frac{1}{2}v_2T_{234} + M(n_2))$ | $\theta_{r,2} \frac{2\gamma_2}{1-e^{-2\gamma_2}}(\frac{1}{2}v_2T_{234} + G(n_2, \gamma_2))$ |
| | Polymorphism $P_{2 \sim (1,3,4)}$ | $\theta_{s,2}L(n_2)$ | $\theta_{r,2} \frac{2\gamma_2}{1-e^{-2\gamma_2}}F(n_2, \gamma_2)$ |
| $3 \sim (1, 2, 4)$ | Divergence $D_{3 \sim (1,2,4)}$ | $\theta_{s,3}(\frac{1}{2}v_3T_{34} + M(n_3))$ | $\theta_{r,3} \frac{2\gamma_3}{1-e^{-2\gamma_3}}(\frac{1}{2}v_3T_{34} + G(n_3, \gamma_3))$ |
| | Polymorphism $P_{3 \sim (1,2,4)}$ | $\theta_{s,3}L(n_3)$ | $\theta_{r,3} \frac{2\gamma_3}{1-e^{-2\gamma_3}}F(n_3, \gamma_3)$ |
| $4 \sim (1, 2, 3)$ | Divergence $D_{4 \sim (1,2,3)}$ | $\theta_{s,4}(\frac{1}{2}v_4T_{34} + M(n_4))$ | $\theta_{r,4} \frac{2\gamma_4}{1-e^{-2\gamma_4}}(\frac{1}{2}v_4T_{34} + G(n_4, \gamma_4))$ |
| | Polymorphism $P_{4 \sim (1,2,3)}$ | $\theta_{s,4}L(n_4)$ | $\theta_{s,4} \frac{2\gamma_4}{1-e^{-2\gamma_4}}F(n_4, \gamma_4)$ |
| $(3, 4) \sim (1, 2)$ | Divergence $D_{(3,4) \sim (1,2)}$ | $\theta_{s,5}(\frac{1}{2}v_5(T_{234} - T_{34}) + J(n_3, n_4, N_3, N_4, T_{34}))$ | $\theta_{s,5} \frac{2\gamma_5}{1-e^{-2\gamma_5}}(\frac{1}{2}v_5(T_{234} - T_{34}) + K(n_3, n_4, \gamma_3, \gamma_4, \gamma_5))$ |
| | Polymorphism $P_{(3,4) \sim (1,2)}$ | $\theta_{s,5}H(n_3, n_4, N_3, N_4, T_{34})$ | $\theta_{r,5} \frac{2\gamma_5}{1-e^{-2\gamma_5}}I(n_3, n_4, \gamma_3, \gamma_4, \gamma_5)$ |

We applied our method to the HDMK tables of the 17, 800 genes. After 200, 000 burn-in steps, we then ran the Markov chain Monte Carlo process for 200, 000 steps to achieve the posterior distributions of the parameters (see Appendix 2 for details). The maximum a posterior (MAP) estimate of the divergence time between orangurans and the common ancestor of humans, chimpanzees and gorillas (*T*_1234_) is 11.6444 (posterior interval: (11.4125, 11.9126)), between gorillas and the common ancestor of humans and chimpanzees (*T*_234_) is 4.9340 (4.6461, 5.2420), between humans and chimpanzees (*T*_34_) is 3.7038 (3.6529, 3.7784), with all values in units of 2*N*_1_. The MAP estimates of the effective population sizes of gorillas, chimpanzees and humans are: *N*_2_ = 0.9665 (0.9458, 0.9878), *N*_3_ = 1.1610 (1.1351, 1.1877), and *N*_4_ = 0.8306 (0.8118, 0.8501), with all values in units of *N*_1_. Our estimates of the demographic parameters are consistent overall with previous studies (Prado-Martinez et al., 2013).

By using orangutans and gorillas as outgroups, we identified 27, 144 fixed synonymous sites and 19, 123 fixed non-synonymous sites in the human lineage (*D*_4∼(1,2,3)_ in Table 2). The average genomic synonymous and non-synonymous divergence are 4.5197 × 10^−4^ and 3.1842 × 10^−4^ (per nucleotide site). We also identified 22, 926 synonymous and 21, 243 nonsynonymous segregating sites in humans (*P*_4∼(1,2,3)_ in Table 2). The average synonymous and non-synonymous densities (per nucleotide site) are 3.8174 × 10^−4^ and 3.5372 × 10^−4^. The ratio of non-synonymous to synonymous divergence sites is smaller than the ratio of non-synonymous to synonymous polymorphisms sites, which is consistent with the fact that the majority of amino acid variations in the genome are deleterious.

### Genes under selection in the human lineage

The histogram in Figure 3(A) shows the posterior distribution of the selection intensity *γ*_4_ for genes under selection in the human lineage. Among these genes, 1211 genes with a 95% confidence interval above 0 were identified as targets under positive selection and 1349 genes with a 95% confidence interval (CI) below 0 are under negative selection. Genes under positive selection in the human-lineage should attract particular attention because they potentially confer to the emergence of human-specific phenotypes and functionality. In the following sections, we focused our analysis on genes showing evidence of positive selection.

**Figure 3:**
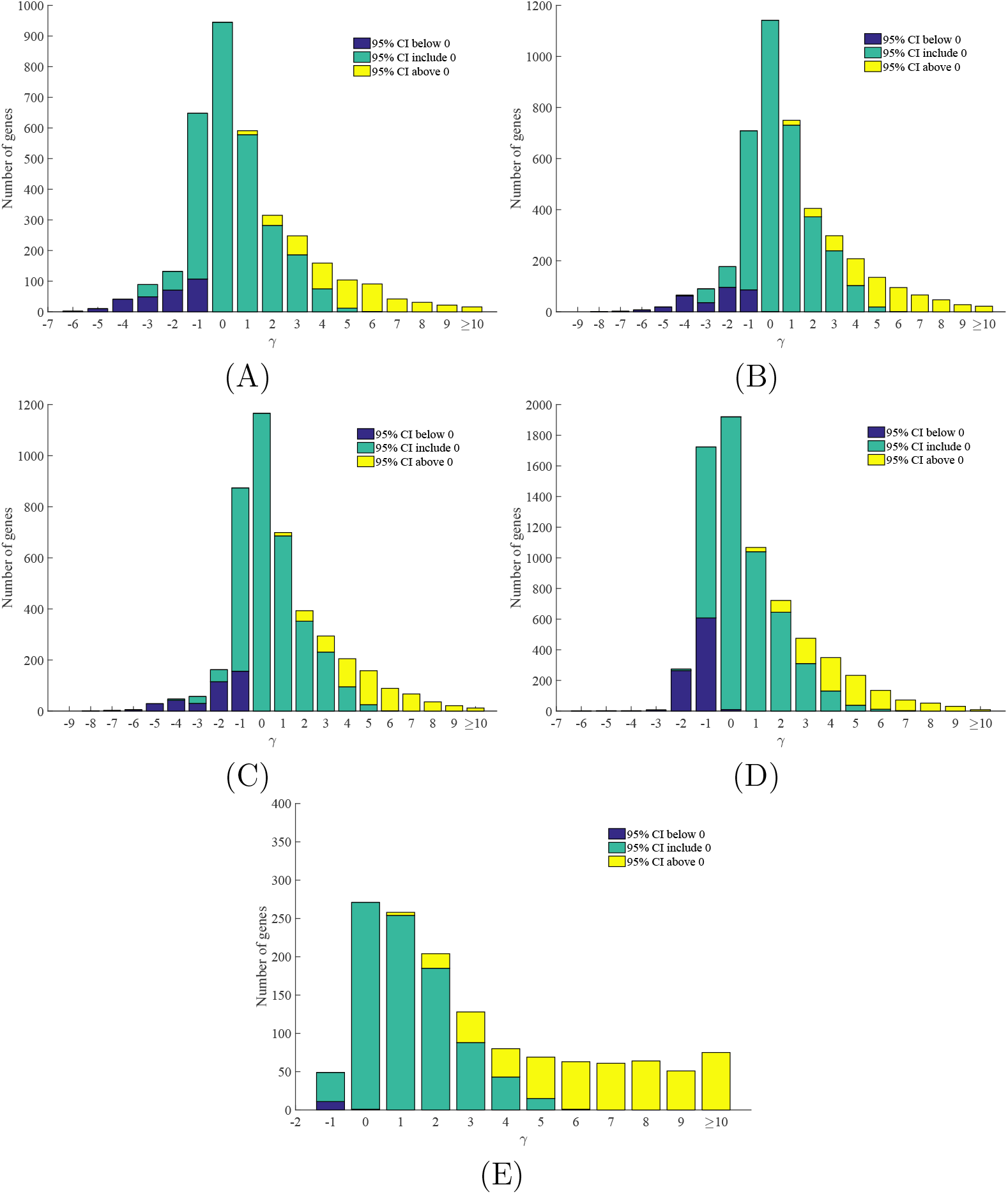
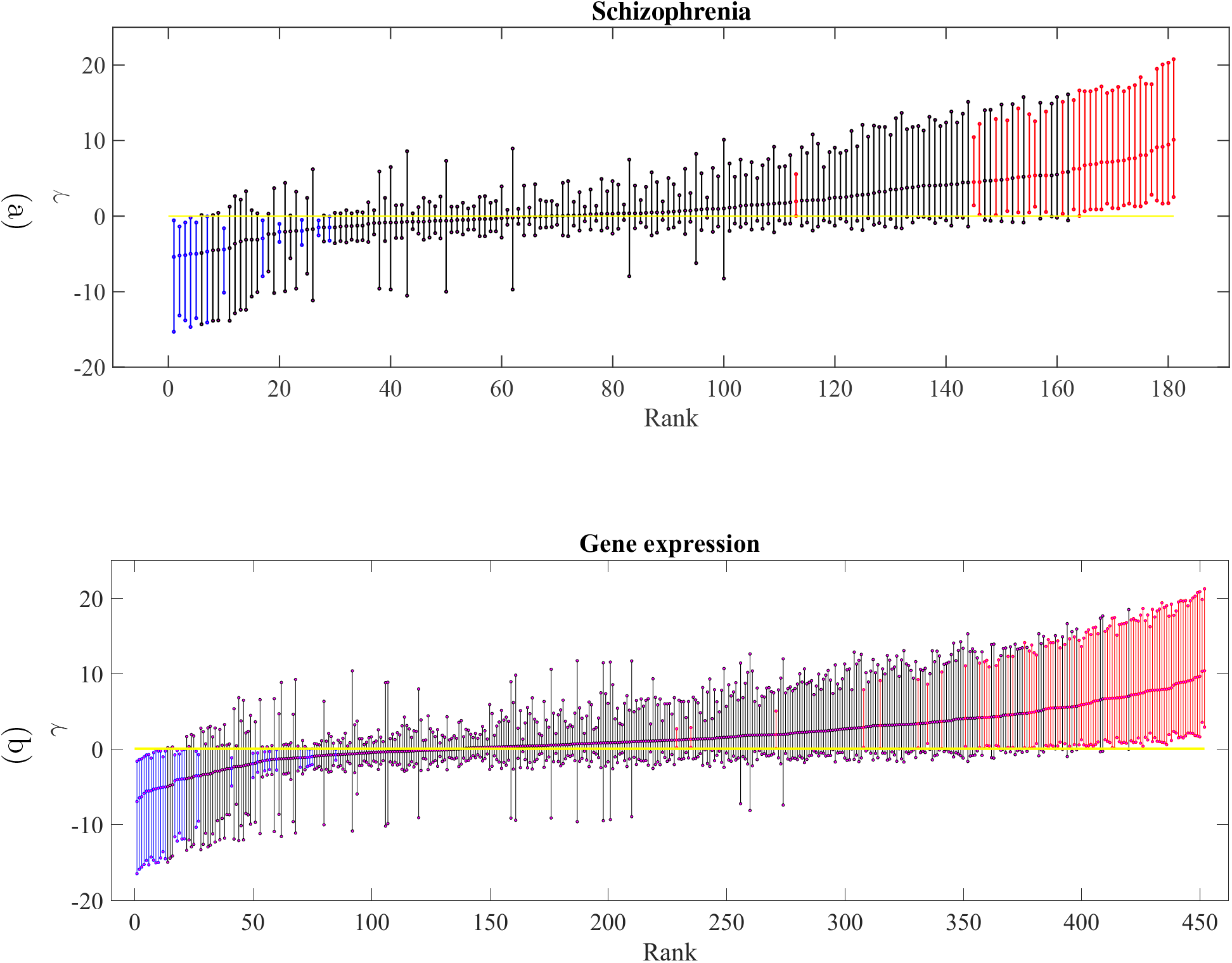
Posterior distribution of *γ* for humans and chimpanzees.

We first investigated the biological functions of those genes under positive selection by performing gene enrichment analyses with the package DAVID (Huang et al., 2009). Gene sets were defined using the KOBAS pathway and disease categories along with Gene Ontology (GO) categories (see Table 4).

Several top pathways in the human lineage are related to the immune systems (*P* < 2.16 × 10^−12^, Table 4), which is similar to the findings of previous studies (Bustamante et al., 2005; Clark et al., 2003; Nielsen et al., 2005). Several genes, such as NCR1, are among the top genes with the highest selection intensity (Table 3): *γ_NCR1_* = 10.07 (2.88, 20.18). Interestingly, these genes are under strong negative selection in the chimpanzee lineage and the human-chimpanzee common ancestor lineage (HC lineage), indicating that accelerated evolution of these genes is likely caused by human-specific resistance to pathogens. These genes belong to different parts of the immune system, i.e., CASP10, DEFB110, HSP90AA1, and INSR, and others are found in the innate immune system; NCR1, UBE2CBP, TRAIP, and SEC24C and other 51 genes are from the adaptive immune system; and another 14 genes, including SAMHD1, NUP107, and UBA7, are related to interferon signaling (see Table 4).

**Table 3:**
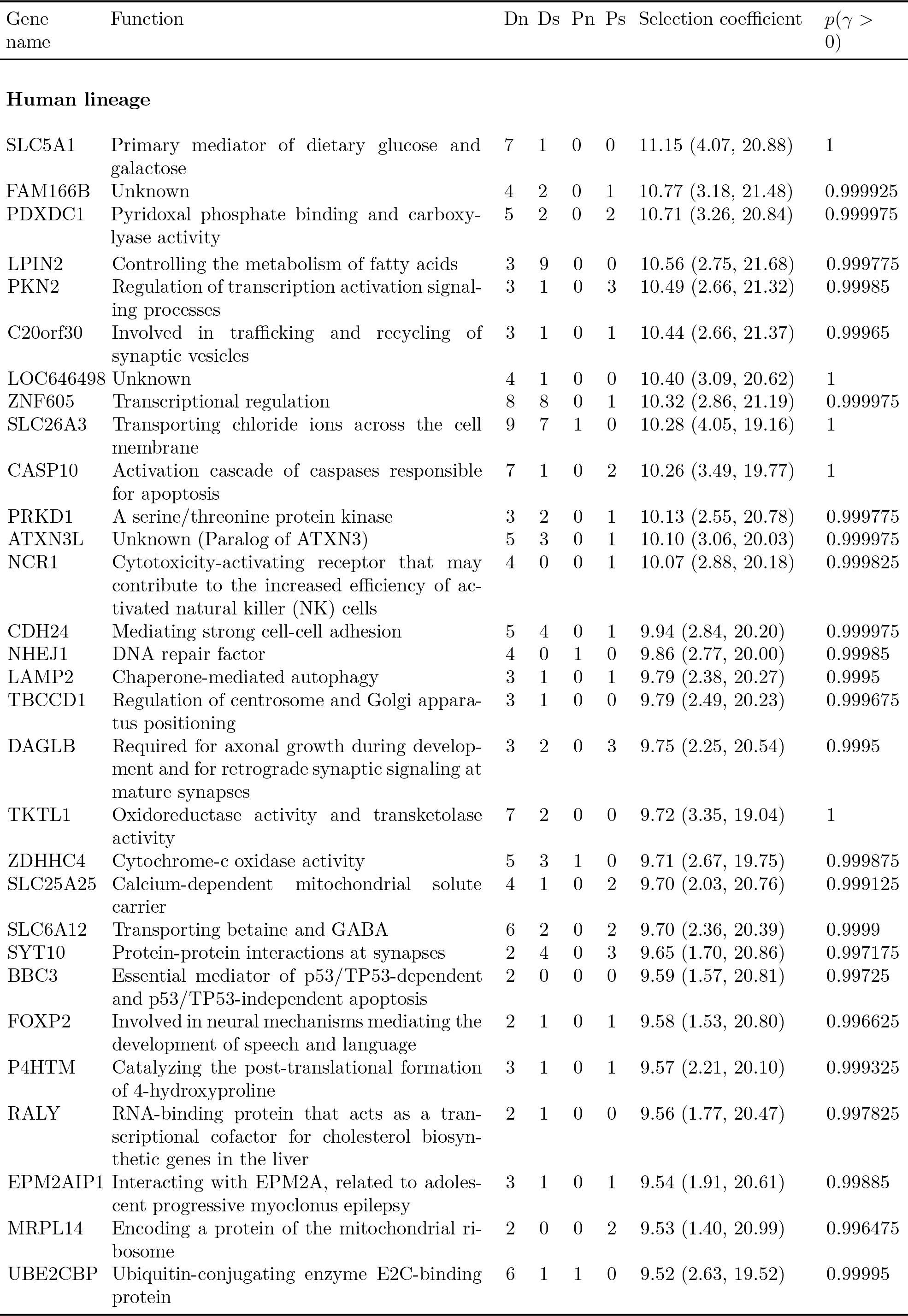
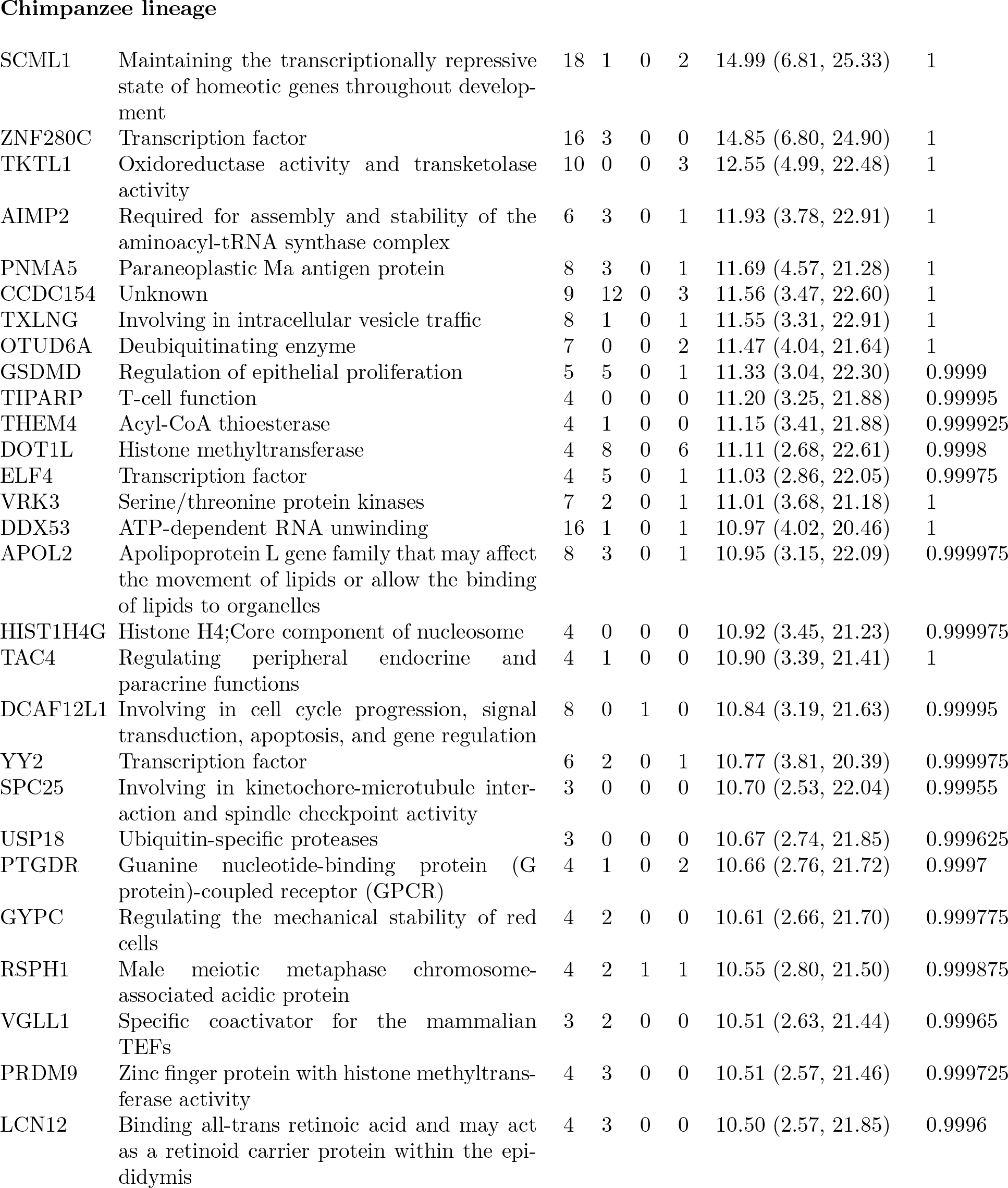
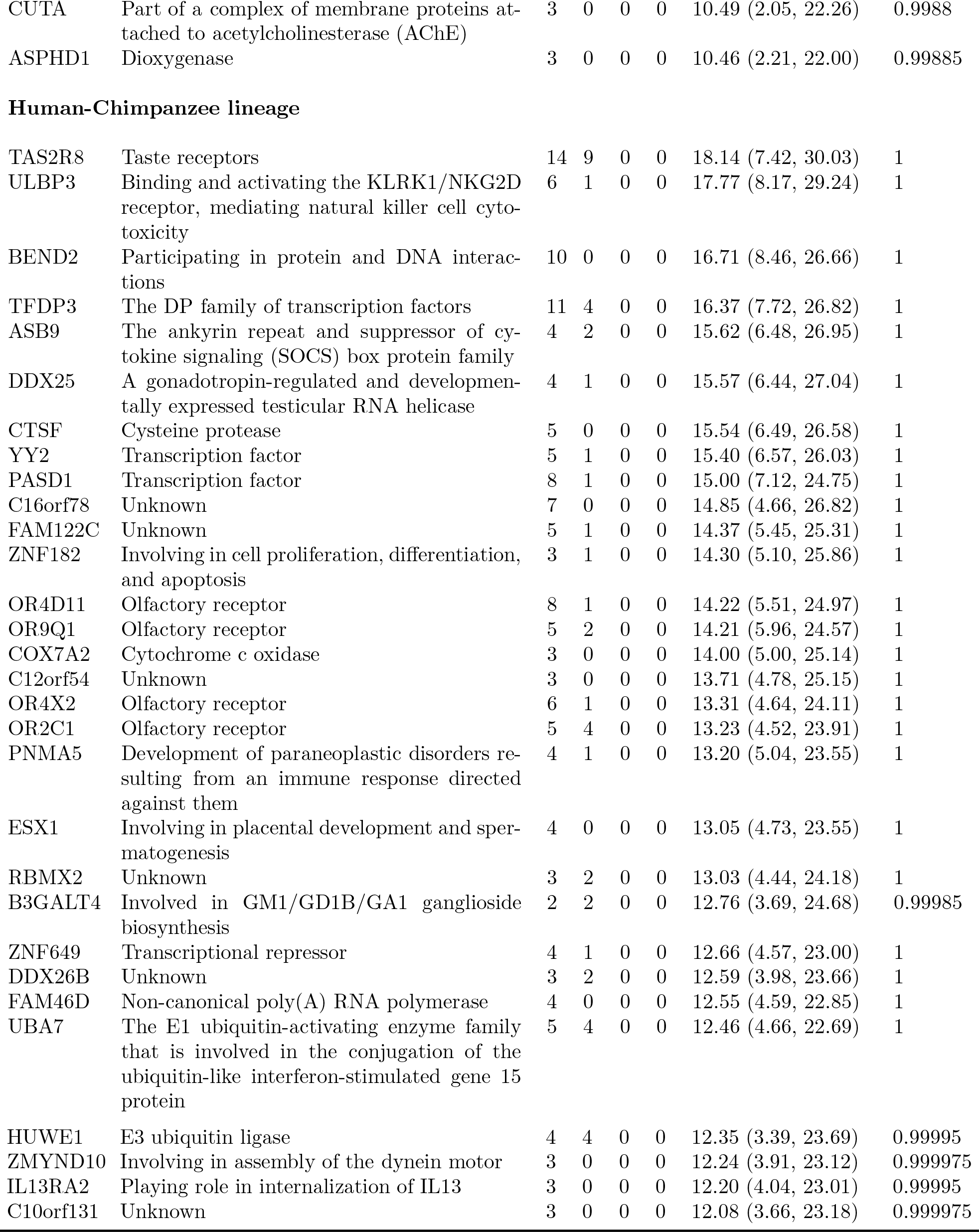
Top 30 positively selected genes in the three lineages

**Table 4:**
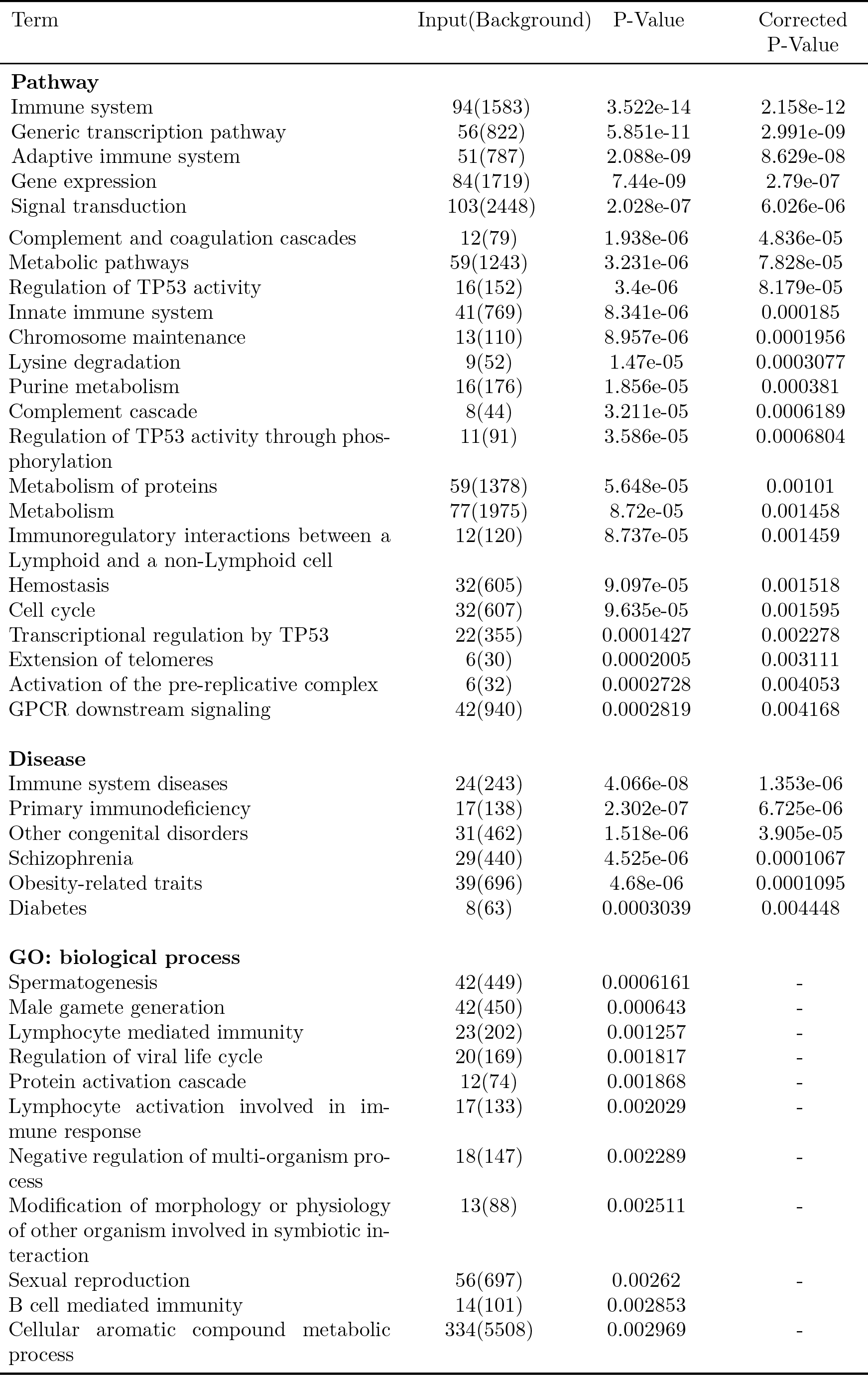
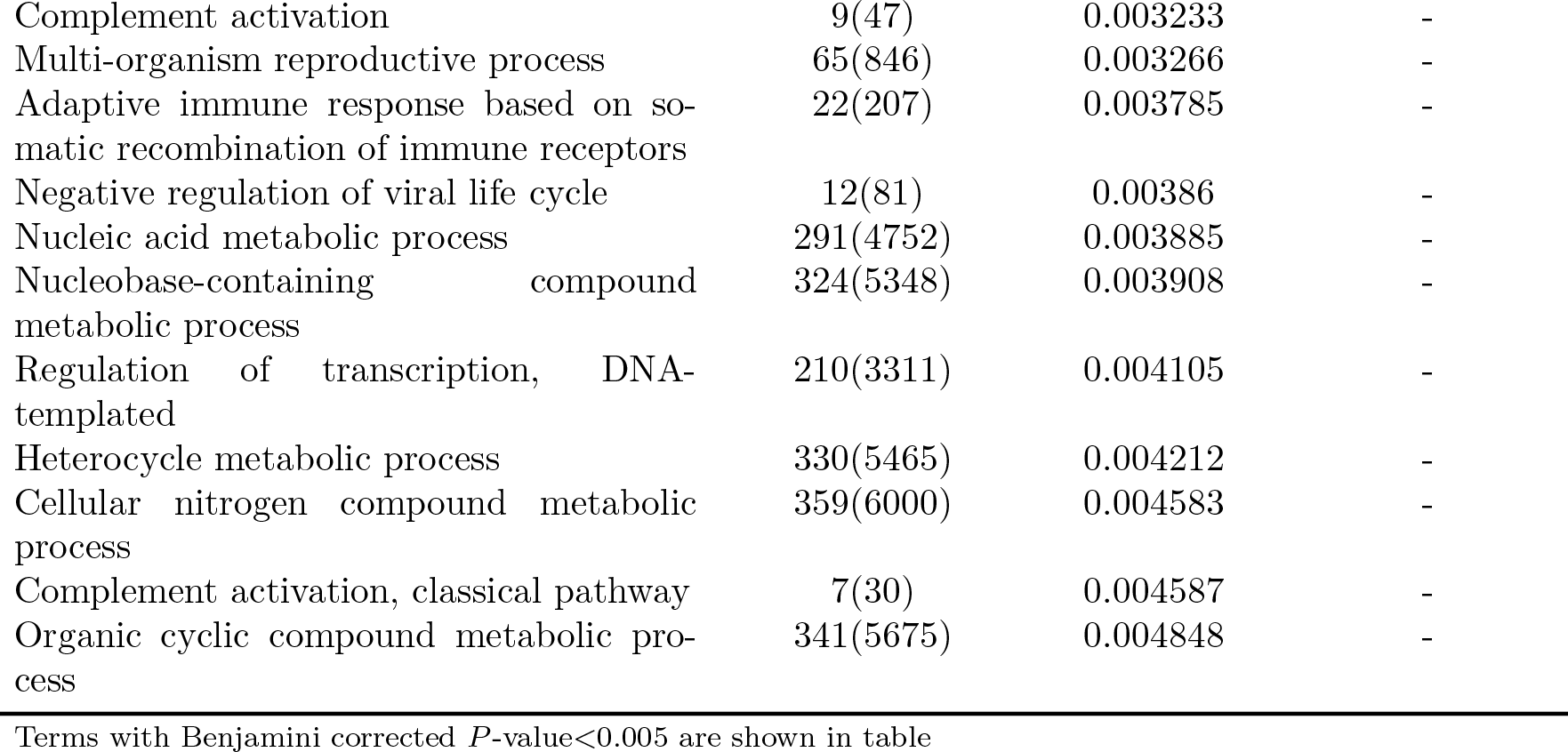
Kobas enrichment of positively selected genes in the Human lineage

| Term | Input(Background) | P-Value | Corrected P-Value |
| --- | --- | --- | --- |
| **Pathway** |  |  |  |
| Immune system | 94(1583) | 3.522e-14 | 2.158e-12 |
| Generic transcription pathway | 56(822) | 5.851e-11 | 2.991e-09 |
| Adaptive immune system | 51(787) | 2.088e-09 | 8.629e-08 |
| Gene expression | 84(1719) | 7.44e-09 | 2.79e-07 |
| Signal transduction | 103(2448) | 2.028e-07 | 6.026e-06 |

| Term | Input(Background) | P-Value | Corrected P-Value |
| --- | --- | --- | --- |
| Complement and coagulation cascades | 12(79) | 1.938e-06 | 4.836e-05 |
| Metabolic pathways | 59(1243) | 3.231e-06 | 7.828e-05 |
| Regulation of TP53 activity | 16(152) | 3.4e-06 | 8.179e-05 |
| Innate immune system | 41(769) | 8.341e-06 | 0.000185 |
| Chromosome maintenance | 13(110) | 8.957e-06 | 0.0001956 |
| Lysine degradation | 9(52) | 1.47e-05 | 0.0003077 |
| Purine metabolism | 16(176) | 1.856e-05 | 0.000381 |
| Complement cascade | 8(44) | 3.211e-05 | 0.0006189 |
| Regulation of TP53 activity through phosphorylation | 11(91) | 3.586e-05 | 0.0006804 |
| Metabolism of proteins | 59(1378) | 5.648e-05 | 0.00101 |
| Metabolism | 77(1975) | 8.72e-05 | 0.001458 |
| Immunoregulatory interactions between a Lymphoid and a non-Lymphoid cell | 12(120) | 8.737e-05 | 0.001459 |
| Hemostasis | 32(605) | 9.097e-05 | 0.001518 |
| Cell cycle | 32(607) | 9.635e-05 | 0.001595 |
| Transcriptional regulation by TP53 | 22(355) | 0.0001427 | 0.002278 |
| Extension of telomeres | 6(30) | 0.0002005 | 0.003111 |
| Activation of the pre-replicative complex | 6(32) | 0.0002728 | 0.004053 |
| GPCR downstream signaling | 42(940) | 0.0002819 | 0.004168 |
| **Disease** |  |  |  |
| Immune system diseases | 24(243) | 4.066e-08 | 1.353e-06 |
| Primary immunodeficiency | 17(138) | 2.302e-07 | 6.725e-06 |
| Other congenital disorders | 31(462) | 1.518e-06 | 3.905e-05 |
| Schizophrenia | 29(440) | 4.525e-06 | 0.0001067 |
| Obesity-related traits | 39(696) | 4.68e-06 | 0.0001095 |
| Diabetes | 8(63) | 0.0003039 | 0.004448 |
| **GO: biological process** |  |  |  |
| Spermatogenesis | 42(449) | 0.0006161 | - |
| Male gamete generation | 42(450) | 0.000643 | - |
| Lymphocyte mediated immunity | 23(202) | 0.001257 | - |
| Regulation of viral life cycle | 20(169) | 0.001817 | - |
| Protein activation cascade | 12(74) | 0.001868 | - |
| Lymphocyte activation involved in immune response | 17(133) | 0.002029 | - |
| Negative regulation of multi-organism process | 18(147) | 0.002289 | - |
| Modification of morphology or physiology of other organism involved in symbiotic interaction | 13(88) | 0.002511 | - |
| Sexual reproduction | 56(697) | 0.00262 | - |
| B cell mediated immunity | 14(101) | 0.002853 | - |
| Cellular aromatic compound metabolic process | 334(5508) | 0.002969 | - |
| Complement activation | 9(47) | 0.003233 | - |
| Multi-organism reproductive process | 65(846) | 0.003266 | - |
| Adaptive immune response based on somatic recombination of immune receptors | 22(207) | 0.003785 | - |
| Negative regulation of viral life cycle | 12(81) | 0.00386 | - |
| Nucleic acid metabolic process | 291(4752) | 0.003885 | - |
| Nucleobase-containing compound metabolic process | 324(5348) | 0.003908 | - |
| Regulation of transcription, DNA-templated | 210(3311) | 0.004105 | - |
| Heterocycle metabolic process | 330(5465) | 0.004212 | - |
| Cellular nitrogen compound metabolic process | 359(6000) | 0.004583 | - |
| Complement activation, classical pathway | 7(30) | 0.004587 | - |
| Organic cyclic compound metabolic process | 341(5675) | 0.004848 | - |

Terms with Benjamini corrected $P$ -value $<0.005$ are shown in table

Another extremely significantly enriched pathway is gene expression (*P* < 2.79 × 10^−7^). This gene set includes 84 genes that fall into four main categories:

(1) RNA polymerase II transcription. This category includes many zinc-finger genes, including ZNF549, ZNF510, ZNF500, ZNF684, and ZNF750, along with others. (2) Transcriptional regulation by small RNAs, including NUP88, NUP133 and NUP107. (3) Regulation of TP53 activity, including AURKB, DNA2, TPX2, RMI1, RBBP8 (through phosphorylation), KAT6A, and EHMT1 (through acetylation or methylation). (4) Regulation of histone methylation, including PHF1, BCOR, CTCFL, MLL2, and BRCA1. Among these genes, JARID2, MLL, and BRCA1 are only under strong lineage-specific positive selection in humans.

The large number of genes from gene expression regulation pathways under positive selection indicates that evolutionary changes at the level of gene regulation played an essential role in the divergence of humans and chimpanzees, which is consistent with the hypothesis of King and Wilson (1975). The evolutionary roles of these genes in forming human-specific phenotypes have yet to be properly recognized, and our results could provide inspiration for further investigations.

Our study also identified genes involved in spermatogenesis under positive selection (Nielsen et al., 2005). High selection intensities were inferred according to the MK table patterns of these genes, including SLC26A3, SEPT4, PRM1, NPHP1 and OVOL1. Among these genes, PRM1 is known for its potential influence on sperm morphology and ability to fertilize eggs (Rooney and Zhang, 1999; Wyckoff et al., 2000; Nielsen et al., 2005).

Metabolism is another large gene category under selection (*P* < 7.83 × 10^−5^) and includes the metabolism of proteins, lipids and lipoproteins, purines, and carbohydrates. Certain genes, such as SLC5A1, show an extremely strong selection signal (*γ* = 11.15 (4.07, 20.88)) in humans, with 7 nonsynomymous divergence sites and 1 synonymous divergence site combined with zero within-species polymorphism sites. This gene is a sodium/glucose cotransporter that mediates the transport of glucose and related substances across cellular membranes, and it is involved in the absorption of glucose. Interestingly, Pontremoli et al. (2015) identified positive selection on the regulatory elements of SLC2A5 and SLCA2, which are fructose or glucose transporters and functionally similar to SLC5A1. However, their coding regions are under neutrality or negative selection in our analysis, which implies that the evolution of carbohydrate consumption pathways may involve both coding regions and regulatory regions. Strong selection on these genes may reflect the importance of starch metabolism and dietary transitions during human evolution. In addition, other genes, such as PKN2, are involved in pathways regulating glucose and lipid metabolism in different tissues, such as skeletal muscle (Ruby et al., 2017).

We also analyze the association between positively selected genes and common diseases (Table 4). Immune system diseases and metabolic diseases, such as diabetes and obesity, are significantly enriched for genes under positive selection in the human lineage. This finding is consistent with the pathway enrichment results. Interestingly, our findings indicate that positive selection of certain genes is correlated with the development of schizophrenia (corrected P = 0.00011). The adaptive evolution of human cognitive abilities was hypothesized to be achieved by pushing the limits of metabolic capabilities, which created side effect such as schizophrenia (Khaitovich et al., 2008).

We then investigated the posterior mean and CIs of genes in these significantly enriched gene categories in further detail. In Figure 4, we present the results for four pathways showing distinct patterns. The genes in each pathway were ranked according to their posterior mean of the selection intensity, with bars representing the 95% CIs. Red and blue colors denote CIs completely above or below the line *γ* = 0, respectively, which correspond to the identification of being positively or negatively selected. Pathways for gene expression and schizophrenia are over-dominantly enriched for positively selected genes. These pathways are putatively under accelerated evolution and thus have played essential roles in the emergence of new functionality. In contrast, certain pathways, such as olfactory signaling, are enriched for negative selection, indicating their conservative evolution in the human lineage. Other pathways, such as metabolism, are significantly enriched for both positive selection and negative selection, thereby showing manifold functionality in evolution.

**Figure 4:**
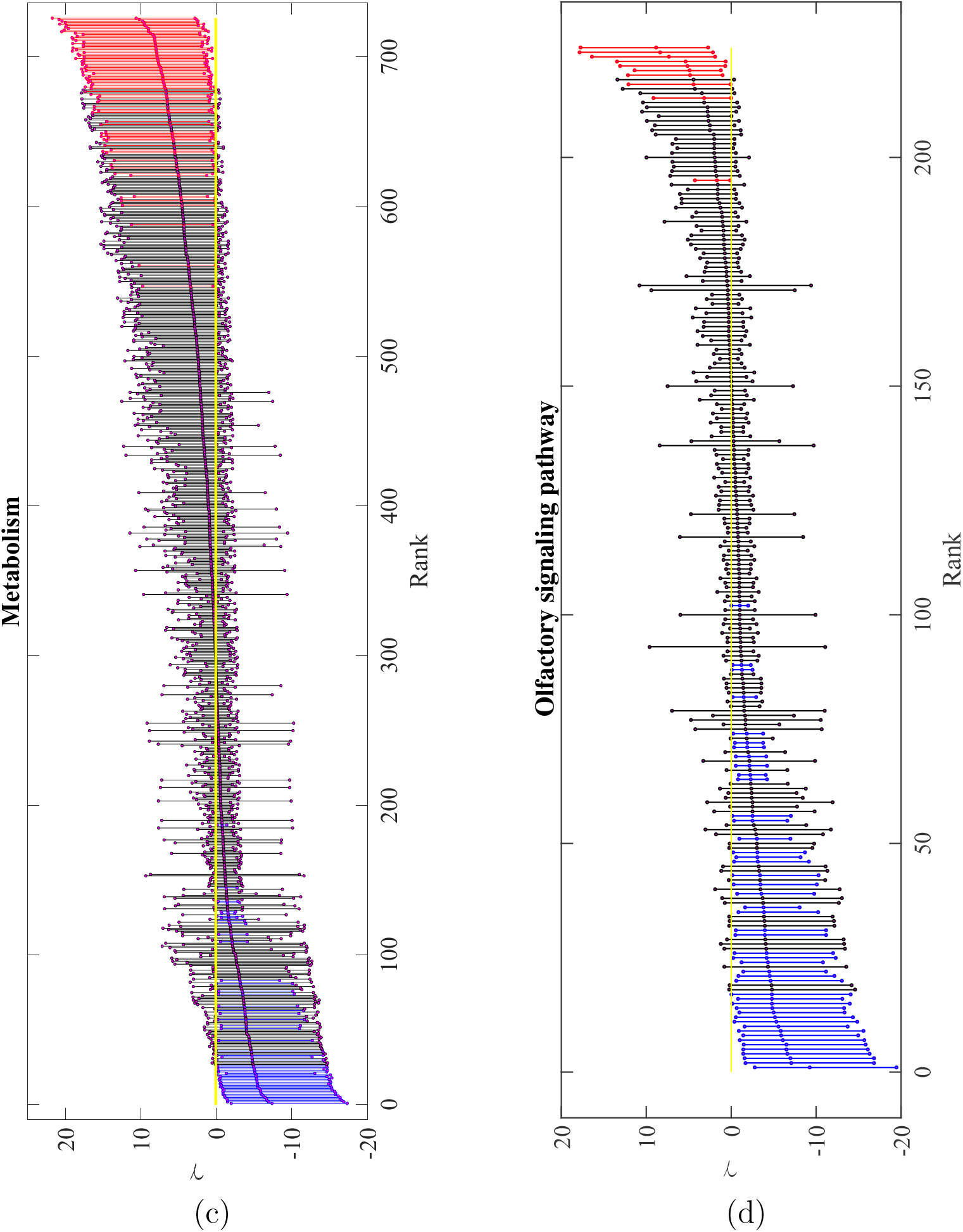
Distribution of y for four pathways or disease-related gene sets.

In addition to performing a categorical analysis of pathways and genes, we also investigated the top 30 genes under positive selection in humans, chimpanzees, and the common ancestor of humans and chimpanzees (see Table 3). The genes were ranked according to their MAP selection intensity. All positively and negatively selected genes are also shown in Figures 5 and 6 as a selection map of the human and chimpanzee genomes. The genes identified as targets of positive or negative selection are marked on each chromosome according to their physical positions and labelled with red and blue, respectively.

**Figure 5:**
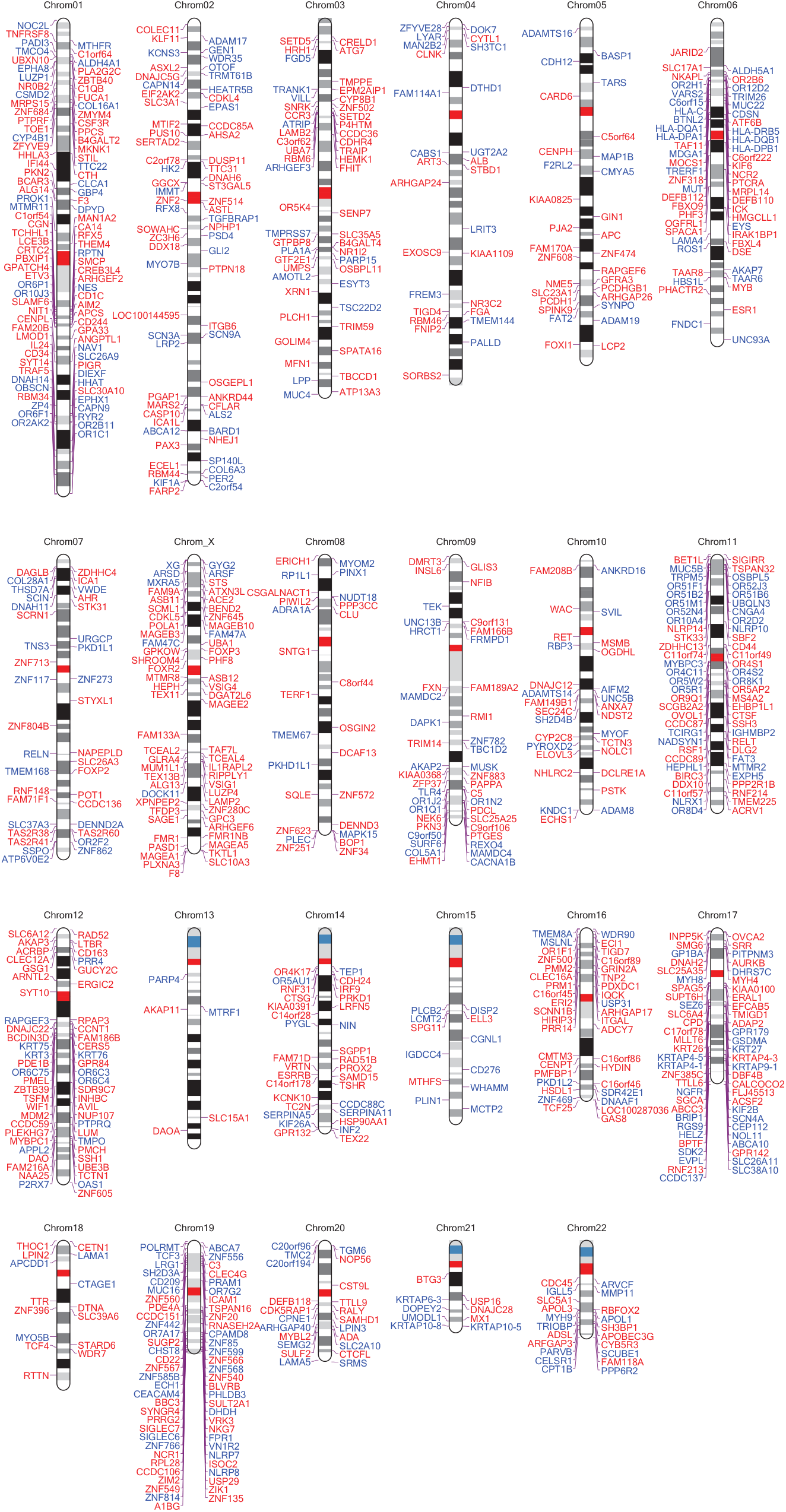
A map of Darwin selection of the human genome.

**Figure 6:**
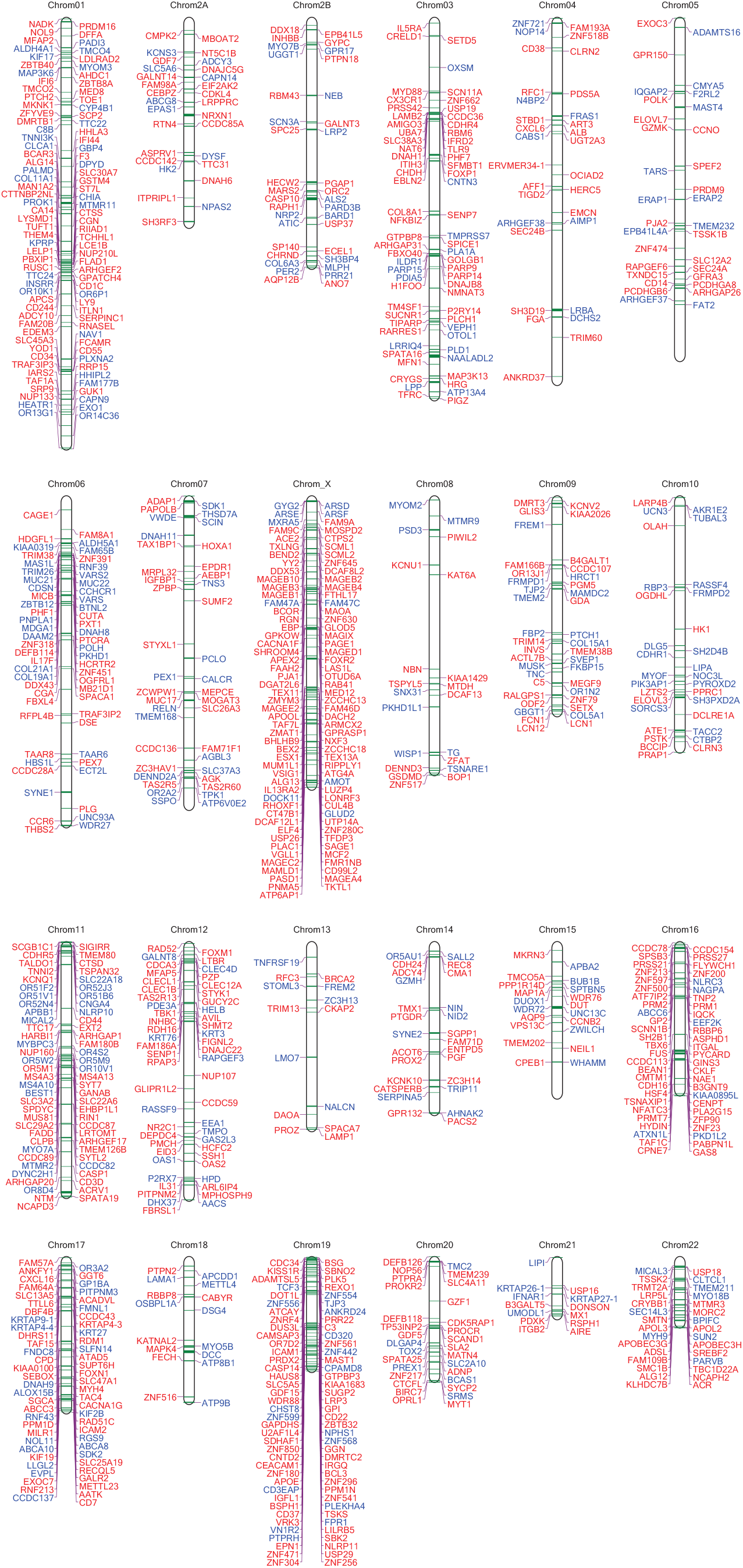
A map of Darwin selection of the chimpanzee genome.

Among the genes from the top list, FOXP2 is well known for its putative functionality in the evolution of language (Enard et al., 2002). Our analysis demonstrates that FOXP2 has been under strong positive selection in humans since the divergence of humans and chimpanzees and presents a selection intensity of *γ* = 9.58 (95% CI. (1.53, 20.80)). FOXP2 is quite conservative in chimpanzees (*γ* = −2.70 (−16.49, 11.43)) and the common ancestor of humans and chimpanzees (*γ* = –1.31 (−16.13, 13.41), which indicates that positive selection on FOXP2 contributes to the emergence of human-specific phenotypes and functionality.

Interestingly, a number of genes on the list are related to neurological systems. RBFOX2 is a conserved RNA binding protein serving as a key regulator of alternative splicing in the nervous system and may play an important role in neuromuscular functions (Raj and Blencowe, 2015). SYT10 is found in pathways of protein-protein interactions at synapses and transmission across chemical synapses. This gene was identified to be important for epileptogenesis in previous studies (Glavan et al., 2009; Woitecki et al., 2016). ATXN3L is a paralog of ATXN3 that has undergone limited study, although GO annotations related to the gene include ubiquitin protein ligase binding and obsolete ubiquitin thiolesterase activity. Mutations of this gene were identified to be correlated with neurodegenerative disease (do Carmo Costa and Paulson, 2012). SLC6A12 is a transporter of betaine and GABA, which may have a role in the regulation of GABAergic transmission in the brain through the reuptake of GABA into presynaptic terminals. EPM2AIP1 interacts with EPM2A, which produces a protein called laforin, and is related to epilepsy (Lafora disease).

### Genes under selection in the chimpanzee lineage

Among the top 30 signals under strong positive selection in chimpanzees (Table 3), SCML1 is the one with the highest inferred selection intensity at 14.99 (6.81, 25.33). SCML1 was identified as a target of repeated positive selection by Nielsen et al. (2005) because it has 15 nonsynonymous and one synonymous substitutions between humans and chimpanzees, although zero polymorphisms are observed in humans. SCML1 is an expression repressor of the Hox genes and important for the developmental differences between humans and chimpanzees. Wu and Su (2008) also identified strong positive selection on SCML1 in multiple primate species and noticed that the gene is expressed in testes, implying its role in testis development and spermatogenesis.

PNMA5 shows a strong signal for positive selection in chimpanzees (*γ* = 11.69 (4.57, 21.28)). The exact molecular function of PNMA5 is still unclear, although previous research indicated that PNMA5 is highly expressed in the neocortex of the brain and may be involved in primate brain evolution. Interestingly, we found that the gene is also under strong positive selection in the human-chimpanzee common ancestor lineage but is under neutral evolution in humans, implying that it once played an important role in the brain evolution of ancient primates but that its effect likely terminated in the human lineage. AIMP2 is another gene known to be related to human neurodegenerative diseases, such as Parkinson’s disease.

### Genes under selection in the common ancestor of humans and chimpanzees

TAS2R8 shows the strongest positive selection signal in the common ancestor of humans and chimpanzees, although it is under strong negative selection in human and under neutrality in chimpanzee (*γHC* = 18.142 (7.4216, 30.0282); *γhuman* = −4.32549 (−15.0859, 2.04698); and *γchimp* = 2.13833 (−0.79666, 7.64481)). TAS2R8 is one of the bitter taste receptor genes and may reflect a change in diet or toxin avoidance during different stages of primate evolution.

Several genes from the top list are olfactory receptors, such as OR4D11, OR9Q1, OR4X2, OR2C1, and OR4F6. Olfactory receptor genes are a large group of genes that show a strong tendency for positive selection (Clark et al., 2003), and our results demonstrate that these olfactory receptor genes have been important for sensory perception ever since the the time of the common ancestor of humans and chimpanzees. Among these genes, some are still actively evolving and under positive selection in humans, such as QR9R1, whereas the rest are conservative and under neutrality or negative selection in humans and/or chimpanzees.

A number of positively selected genes in the common ancestor lineage of humans and chimpanzees are related to transcription regulation, including TFDP3, YY2, PASD1, ZNF182, ESX1, ZNF649, and ZMYND10. In particular, some of these genes are involved in spermatogenesis. ZMYND15, for example, encodes a histone deacetylase-dependent transcriptional repressor that is important for spermatogenesis and male infertility (Yan et al., 2010). ZMYND10 is a zinc finger gene functioning in assembly of the dynein motor, and it is highly expressed in sperm and important for sperm movement.

ASB9, PNMA5, ICAM1 and IL13RA2 are found in the immune system. ICAM1 is a receptor that mediates the binding of *Plasmodium falciparum* to erythrocytes. Mutations on ICAM1 were under very recent positive selection in human populations (Kun et al., 1999). We found that the gene is under strong selection in the common ancestor of human and chimpanzee (*γ* = 9.81 (3.22, 19.27)), which is consistent with the long history of malaria as a pathogen of primates and may reflect the gene’s role in resistance to malaria since the early stages of primate evolution. We also noticed that ICAM1 is under selection in both chimpanzees (*γ* = 8.71(3.15, 16.95) and humans (*γ* = 4.95(1.19, 11.25)). Humans and chimpanzees are affected by different malaria species (*P. falciparum* and *P. reichenowi*, Martin et al. (2005)). ICAM1 is likely undergoing parallel evolution for resistance to the two parasites. Further investigation of the divergence sites among the three species may provide insights into the mechanism of malaria resistance.

Overall, in the above study, we discussed the genes and pathways under strong positive selection in humans, chimpanzees and their common ancestral lineage. Nielsen et al. (2005) used a likelihood-ratio test to compare the dN/dS ratios for coding regions of one human and one chimpanzee genome and provided a list of the 50 top genes under positive selection, and we found that 12 of their genes are also identified as significant in our analysis. However, only 7 out of the 12 genes are under positive selection in the human lineage; three are under positive selection in chimpanzees (CD72, SLC22A4 and DFFA), and the other two genes (RBM23 and C16orf3) are under selection only in the common ancestor of human and chimpanzee. Some other genes, such as SPATA3, are not significantly under positive selection. The MK table entry for SPATA3 in the human lineage is (1, 1, 3, 1), indicating that positive selection on SPATA3 is not likely. The difference between our results and those of Nielsen et al. (2005) demonstrates that by extending the MK-like or dN/dS tests to multiple species, our method can gain information for precisely distinguishing selection that occurred in the human lineage, in the chimpanzee lineage, or in the common ancestor of the two species, thereby providing additional insights into understanding the selection process.

## Discussion

We present the computational method HDMKPRF for detecting lineage-specific selection by jointly analyzing polymorphism and divergence data from multiple species. HDMKPRF is an extension of the MKPRF method and can be used from two species to multiple species (Bustamante et al., 2001). Our method has several advantages over existing approaches. The pairwise comparison of two species in both the MK test and MKPRF is directionless, and it only identifies genes that are divergent between the two species or populations but provides no further conclusions as to the lineage under selection (Akey et al., 2002; Bustamante et al., 2005; Clark et al., 2003; Chen et al., 2010). However, the HDMKPRF method can pinpoint the occurrence of selection to a specific lineage, including the internal lineages. As we demonstrated in the analysis of the Hominidae data, HDMKPRF identifies specific targets of selection in human and chimpanzee lineages as well as the common ancestor of humans and chimpanzees.

Similar to the MK test and MKPRF, the HDMKPRF method analyzes both polymorphism and divergence sites, which increases the power for detecting selection and helps distinguish positive selection from the relaxation of negative selection (Wyckoff et al., 2000). By using the Bayesian approach to combine information from all gene loci, the method is more powerful than single-locus methods, such as the MK test and dN/dS ratio tests. We apply the HDMKPRF method to the genomic sequences of four species of Hominidae and identify gene loci under lineage-specific positive and negative selection. Cagan et al. (2016) recently analyzed the same data set using the MK test and only identified a limited number of genes under positive selection since the divergence of humans and chimapnzees. A comparison of our results with former studies demonstrates that the HDMKPRF method outperforms alternative methods with higher power and provides additional insights into the distribution of selection effects and the temporal and spatial occurrence of Darwinian selection over evolutionary history.

With the development of sequencing technologies, genomic population data for multiple species are abundant, thus necessitating methods that can efficiently analyze both within-species and between-species data. The HDMKPRF method presented in this paper satisfies such a need, and we expect that its application potential will be extensive in comparative genomic studies.

## Appendices

## A1. Segregating site pattern *P*_(2,3)∼1_, *P*_(3,4)∼(1,2)_ and *D*_(3,4)∼(1,2)_

For the three-species scenario, we assume that the allele frequency x of the common ancestor of species 2 and 3 (species 4) at time *T*_23_ follows the stationary distribution under neutrality for synonymous sites (*f* (*x*), Equation 1) and distribution under selection for replacement sites (*g*(*x*), Equation 6). After the population split and with evolution over time, the joint allele frequency distribution of species 2 (*y*) and species 3 (*z*) of synonymous sites is as follows: 
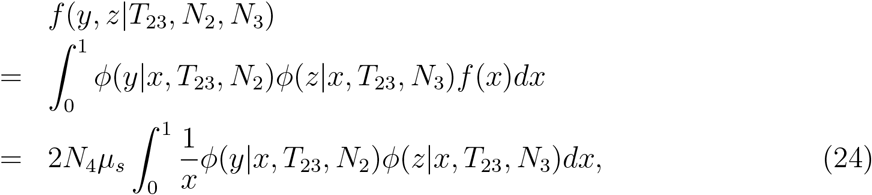
where *ϕ*(*y*|*x*,*T*,*N*) represents the transient allele frequency distribution *y* given its initial frequency *x* at time *T* and population size *N* (Chen et al., 2007). Kimura (1955a) found that for neutral evolution 
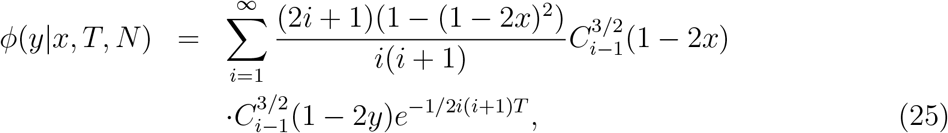
where 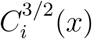 is the Gegenbauer polynomial with λ = 3/2.

Kimura (1955b) provides the transient distribution for alleles under selection, which is in complicated form and is difficult to calculate. Williamson et al. (2005) and Evans et al. (2007) adopted the Crank-Nicolson finite difference method to approximate the transient distribution as the solution of a forward diffusion equation, and several recent studies provided analytical solutions using perturbation methods (Schraiber, 2014; Živković et al., 2015). Similar to the neutral case, the joint allele frequency distribution of nonsynonymous sites under selection in species 2 and 3 is: 
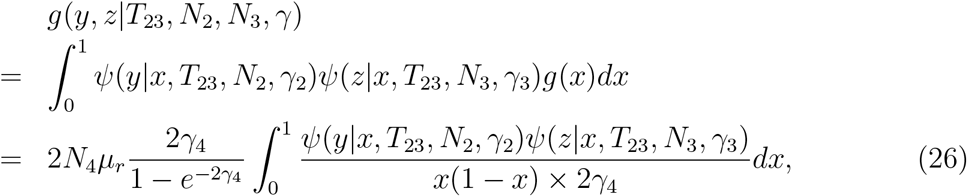
where ψ(*y*|*x*,*T*, *N*, *γ*) represents the transient allele frequency distribution for *y* with an initial allele frequency *x*, time *T*, population size *N* and selection intensity *γ*.

The expected number of polymorphism synonymous sites segregating in species 2 and 3 is 
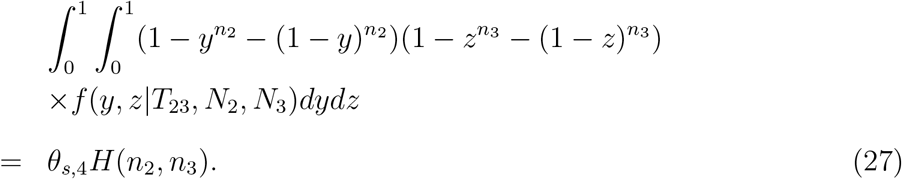

The expected number of polymorphism replacement sites segregating in species 2 and 3 is as follows: 
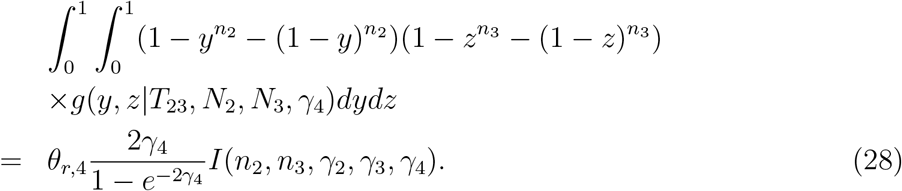

The other entries of the three-species MK table are now different from that of Table 1. For example, the expected number of *P*_2∼(1,3)_ includes two components. The first part consists of sites fixed in sample 3 but still segregating in sample 2: 
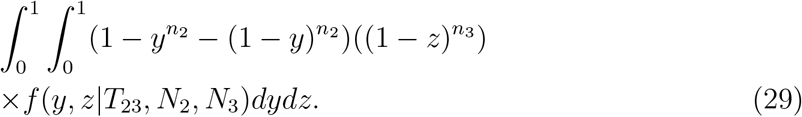

The second part consists of the new mutations occurring in species 2 since *T*_23_: 
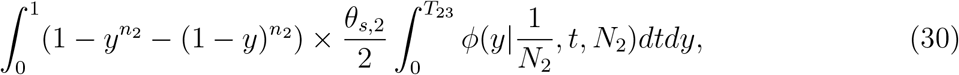
where 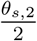 is the number of new neutral mutations that enter the population every generation with the initial frequency of 1/*N*_2_.

Note that the above formula for *P*_*s*,2∼(1,3)_ is different from that in Table 1 and is applicable in different situations. An empirical criterion for choosing the two fomulae is based on the distribution of TMRCA of *n*_2_. According to Griffiths (1984), the TMRCA asymptotically follows a normal distribution, with the mean and variance determined by *N*_2_ and the population history (see Griffiths (1984) and Chen and Chen (2013) for detailed formulae); thus Pr(TMRCA < *T*_23_). When *T*_23_ is sufficiently large and Pr(TMRCA < *T*_23_) > 0.95, we adopt the formulae in Table 1; otherwise, we use Equation 29 and 30.

For the four-species scenario, we can obtain the expected values for *P*_*s*,(3,4)∼(1,2)_ and *P*_*r*,(3,4)∼(1,2)_ via a similar method used for *P*_*s*,(2,3)∼1_ and *P*_*r*,(2,3)∼1_. Synonymous divergence sites *D*_(3,4)∼(1,2)_ in the four-species scenario include two components: the sites fixed on the branch between species 5 and 6 and the sites fixed in the sample of *n*_3_ and *n*_4_. Following Equation 5, the first component is as follows: 
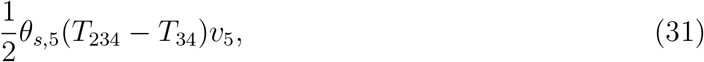
with *υ*_5_ = *N*_1_/*N*_5_. The second component includes those sites fixed in the sample of *n*_3_ and *n*_4_: 
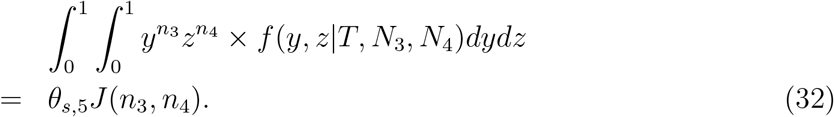

Similarly for replacement sites, the two components are: and 
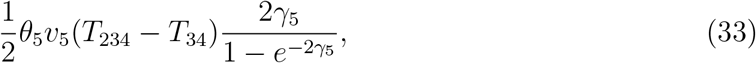
and 
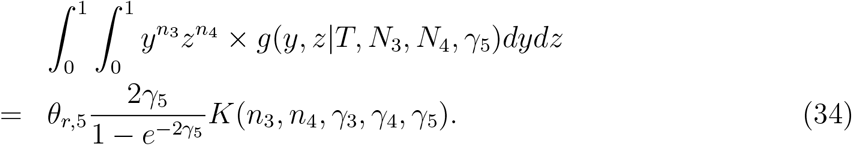

Therefore, we have *P*_(3,4)∼(1,2)_ and *D*_(3,4)∼(1,2)_.

## A2. MCMC steps for parameter optimization for the 4-species McDonald-Kreitman tables

In the Bayesian Poisson random field model of four species, the parameters include 
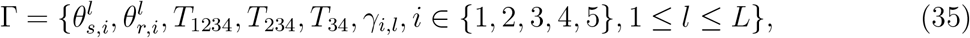
where *i* is the index of lineages (species, see Figure 1B), *l* is the index of genes. *D* and *P* are four-species data containing lineage-specific divergence and polymorphism sites (1 ∼ (2, 3, 4) etc.). Since the posterior distributions of Γ are analytically intractable, we apply the Markov chain Monte Carlo method (MCMC) to achieve them. The details of MCMC steps are as follows.

### Initialization

The initial values of divergence time *T*_1234_, *T*_234_ and *T*_34_ are chosen according to prior knowledge or with arbitrary positive values satisfying *T*_1234_ > *T*_234_ > *T*_34_. The selection parameters *γ_i_*, *i* = 1, 2, 3, 4, 5 are generated from the normal distribution *N*(0, 8). We denote *υ_i_*, *i* = 2, 3, 4, 5 as the ratio of effective population sizes *N*_1_/*N*_i_. In the four-species McDonald-Kreitman table, the number of polymorphism synonymous mutations occurring in gene *l* in lineage *i*, with *i* = 1, 2, 3, 4 (as is shown in Figure 1B), is 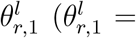. Therefore, for *i* = 2, 3, 4, *υ_i_* can be estimated as follows: 
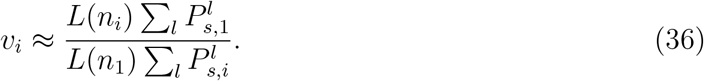

### MCMC iteration

Locus-specific mutation parameters for gene l in lineage 1 ∼ (2, 3, 4) are updated via Gibbs sampling. For each gene *l*, given values of *γ_i,l_*, *i* = 1, 2, 3, 4, 5, a value of 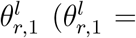 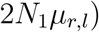 is generated from a gamma distribution 
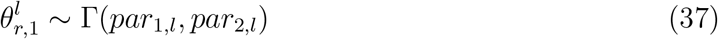
with 
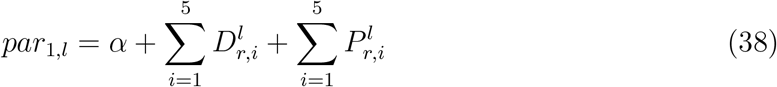
and 
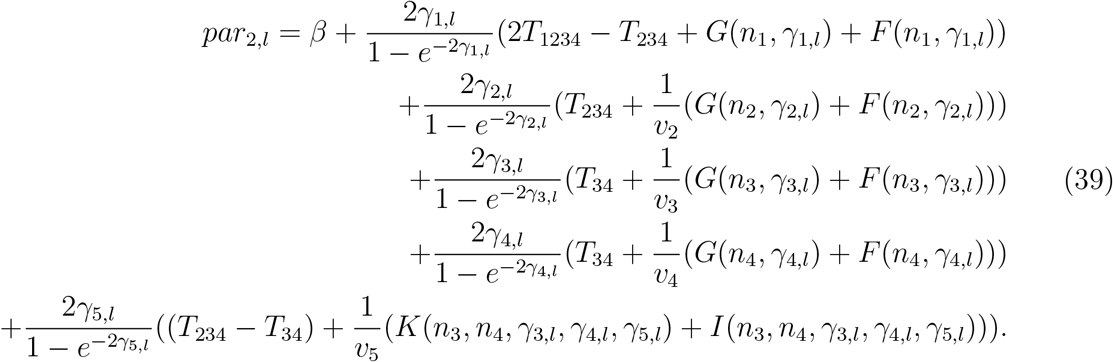
Since *D*_(3,4)∼(1,2)_ and *P*_(3,4)∼(1,2)_ are usually of low information with high volatility in our example, we approximate the posterior distribution of 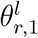 
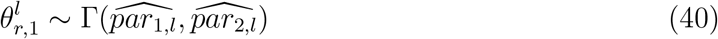
with 
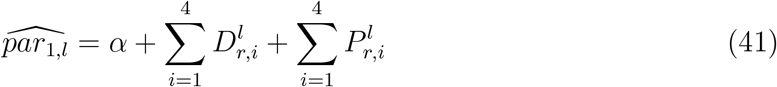
and 
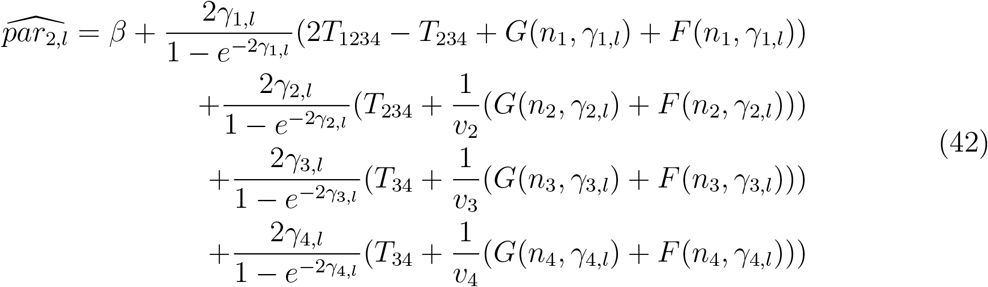

Similarly, a value for 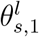 was generated from the gamma distribution 
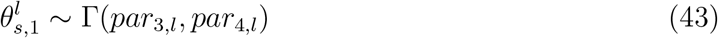
with 
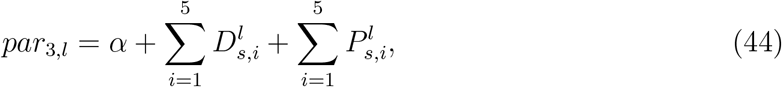
and 
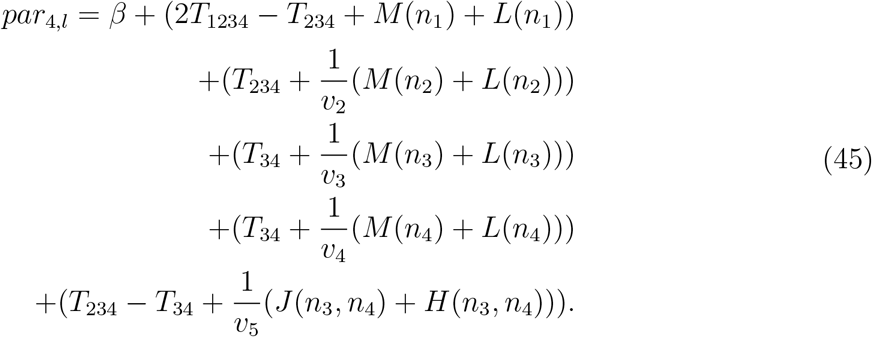

We also approximate the gamma distribution: 
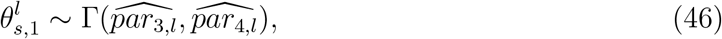
where 
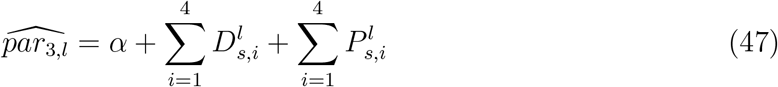
and 
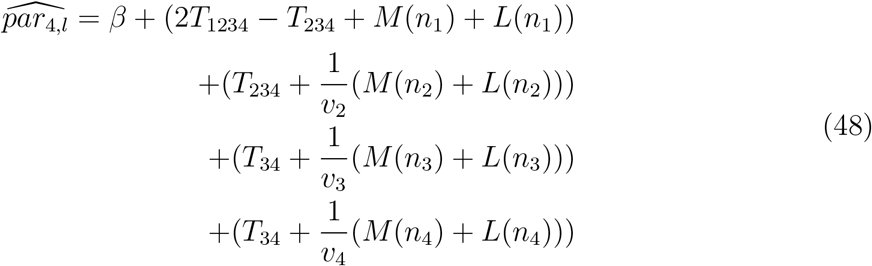

Note that in the above gamma distributions, *α* and *β* are uninformative small values close to 0.

Once 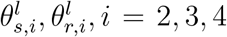 are updated, 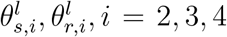 can be calculated by 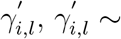 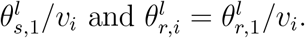

Then, we update the selection parameters using Metropolis sampling as shown in Algorithem 1.

#### Algorithem 1: Updating selection parameters by Metropolis sampling

~~~
For selection coefficient *γ_i,l_* of gene *l* of lineage *i*
        Sampling 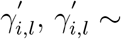 *Uniform*[*γ_i,l_* – *ε, γ_i,l_* + *ε*]
                If 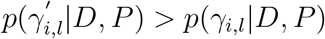
                        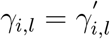
                Else
                        sample *u*, *u* ∼ *Uniform* [0,1]
                        If 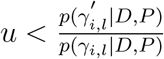
                                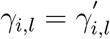
                       Else
                        *γ*_*i*,*l*_ = *γ*_*i*,*l*_
                      End
               End
End
~~~

In Algorithem 1, *p*(*γ_i,l_*|*D,P*) is the posterior probability for *γ_i,l_*. 
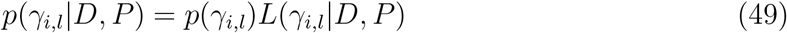
with 
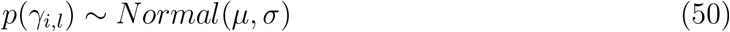
and 
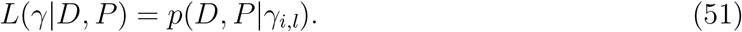

Considering that selection coefficient *γ_i,l_* only affects the non-synonymous mutation numbers, we have 
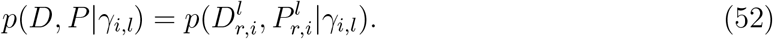

Since the expected *K* and *I* in the four-species McDonald-Kreitman table are difficult to calculate and only have a weak influence on the estimation of *γ_5,l_*, we approximate the likelihood of *γ_5,l_* as follows: 
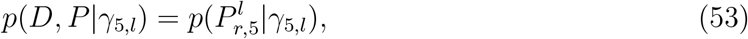

*E* 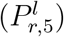 is simplified as follows: 
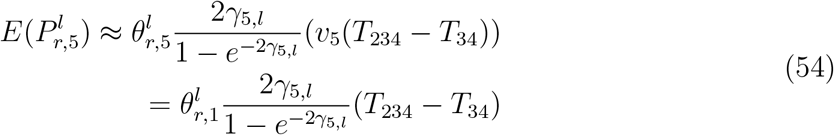

Once all the selection and mutation parameters have been updated, the divergence times *T*_1234_, *T*_234_ and *T*_34_ are updated by Metropolis sampling in a manner analogous to the updating of the γ values. The steps for updating divergence time are shown in Algorithem 2 below.

#### Algorithem 2: Updating divergence time by Metropolis sampling

~~~
For divergence times *T* in {*T*_1234_, *T*_234_, *T*_34_} in turn
        Sampling *T′*, *T′* ∼ *Uniform*[*T* – *ε, T* + *ε*]
        If *p*(*T′*|*D,P*) > *p*(*T*|*D,P*)
                *T* = *T′*
        Else
                Sampling *u,u* ∼ *Uniform*[0,1]
                If 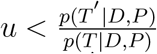
                        *T* = *T′*
                Else
                        *T* = *T*
                End
       End
End
~~~

In Algorithm 2, divergence time *T*_1234_ affects divergence sites *D*_1∼(2,3,4);_ *T*_234_ affects *D*_1∼(2,3,4)_, *D*_2∼(1,3,4)_ and *D*_(3,4)∼(1,2);_ and *T* 34 affects *D*_3∼(1,2,4)_, *D*_4∼(1,2,3)_ and *D*_(3,4)∼(1,2)_: 
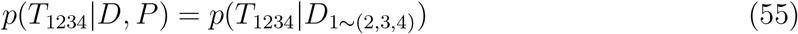

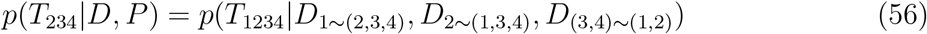

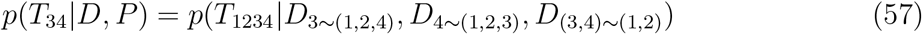

Note that in Algorithm 2, *T* has to satisfy *T*_34_ < *T*_234_ < *T*_1234_. In our algorithm, we only use the synonymous divergence mutant numbers in the MH sampling step to update *T*. The simulation results show that this processing method can improve the performance of divergence time estimation.

The total number of MCMC iterations was 400,000, and the first 200,000 iterations were treated as burn-in and disregarded; subsequently, we sampled data points every 10 iterations, which yielded a total of 40,000 sample points.

## Acknowledgements

This project was supported by the “Strategic Priority Research Program” of the Chinese Academy of Sciences (Grant No. XDB13020400), the National Natural Science Foundation of China (Grant No. 91731302, 31571370 and 91631106), and the “Hundred Talents Program” of the Chinese Academy of Sciences.

